# Assessing Extracellular Vesicle Turnover In vivo Using Highly Sensitive Phosphatidylserine-Binding Reagents

**DOI:** 10.1101/2024.11.14.623541

**Authors:** Lavinia Flaskamp, Monica Prechtl, Annkathrin Scheck, Wenbo Hu, Christine Ried, Georg Kislinger, Mikael Simons, Anne Krug, Jan Kranich, Thomas Brocker

**Affiliations:** Institute for Immunology, Faculty of Medicine, BMC, LMU Munich, Großhaderner Strasse 9, 82152 Planegg; German Center for Neurodegenerative Diseases (DZNE), Munich, Germany

**Keywords:** Extracellular Vesicles (EVs), Phosphatidylserine (PS), MFG-E8, Lactadherin, EV-turnover

## Abstract

Phosphatidylserine (PS) is a well-established marker for apoptotic cells and activated platelets, and it is also found on extracellular vesicles (EVs). However, the frequency of PS^+^ EVs remains a topic of debate, probably due to the distinct lipid composition of EVs, which varies with the source material. This variation underscores the need for highly sensitive detection reagents to accurately differentiate between PS^+^ and PS^−^ EVs. To address this issue, we compared several PS-binding reagents for the staining of blood EVs, including milk fat globule factor E8 (MFG-E8, lactadherin), a derivative of MFG-E8, and Annexin V, the most used compound in the literature. Here, we demonstrate the predominance of PS^+^ EVs in both murine and human blood and reveal significant differences in the affinities of the PS-binding proteins. With approximately 90% of the detected blood EVs being PS^+^, we utilized a derivative of MFG-E8 for *in vivo* monitoring of both circulating and cell-bound EVs, providing a novel tool to analyze endogenous EV kinetics. Our findings showed that murine PS^+^ plasma EVs were rapidly cleared from circulation (∼50% after 30 minutes), yet they remained present on splenic B cells and monocytes/macrophages. In conclusion, MFG-E8-based reagents enable externalized PS to serve as a universal EV marker, greatly improving EV isolation, standardizing EV quantification, and facilitating diagnostic EV monitoring and disease biomarker discovery.

## INTRODUCTION

Over the past two decades, research on EVs has intensified, revealing novel roles for these particles in a wide range of physiological and pathological contexts. Moreover, EVs are being explored as potential cargo-delivery vehicles in various therapeutic applications. Despite this growing interest, the study of sub-micron-sized particles presents inherent challenges. Currently, EVs are primarily characterized and classified based on their size or biogenesis (Couch et al., 2021), as there is no universally accepted marker for EVs. Broadly recognized markers remain scarce or entirely lacking.

PS, a negatively charged phospholipid, is typically localized to the inner leaflet of cellular membranes. However, during apoptosis, PS is externalized. The enzyme flippase plays a crucial role in maintaining PS within the inner leaflet of the plasma membrane, while scramblase facilitates its movement to the outer leaflet. (Daleke, 2003). Loss of membrane asymmetry, resulting from flippase inactivation and scramblase activation, facilitates membrane disintegration and the subsequent release of extracellular vesicles (EVs) (Benedikter et al., 2018; Shao et al., 2018; Stahl et al., 2019). Consequently, PS is exposed on most EVs (Hugel et al., 2005; Llorente et al., 2013; Martinez & Freyssinet, 2001; Skotland et al., 2020; Thery et al., 2002), although it is not universally present across all EV subspecies (Connor et al., 2010). The question of which types and how many EVs are actually PS^+^ remains a topic of ongoing debate. Apoptotic bodies are commonly accepted as PS^+^ EVs due to PS externalization during cell death (Atkin-Smith et al., 2017; Caruso & Poon, 2018). In contrast, the presence of PS on exosomes - another subclass of small Evs - is more controversial, given their origin from multivesicular bodies and the endolysosomal pathway (Kalluri & LeBleu, 2020; Skotland et al., 2017; Tripisciano et al., 2017). Additionally, PS has been identified as a selective marker for EVs derived from specific malignancies (Lea et al., 2017; Sharma et al., 2017), tumor cells (Matsumura et al., 2019; Perez et al., 2023), and pre-eclampsia (Lalic-Cosic et al., 2021). However, it is not considered a marker for all types of EVs.

A recent lipidomics study of EVs revealed distinct lipid compositions and varying levels of PS depending on the source material (Groß et al., 2024). For example, EVs from semen and saliva were found to be rich in PS, while those from blood exhibited very low levels of PS (Groß et al., 2024). Nonetheless, all EVs displayed some degree of PS exposure (Groß et al., 2024), which contrasts with earlier reports suggesting that only tumor-derived EVs were PS^+^ (Lea et al., 2017; Matsumura et al., 2019; Perez et al., 2023; Sharma et al., 2017). Thus, while there is evidence that PS could potentially serve as a universal marker for EVs, a highly sensitive PS-specific reagent is still needed to enable reliable detection across a diverse range of PS concentrations.

Externalized phosphatidylserine (PS) can be detected by various molecules. For instance, milk-fat globule factor E8 (MFG-E8, also known as lactadherin), a protein produced by macrophages and follicular dendritic cells (Kranich et al., 2008), facilitates the phagocytic clearance of PS^+^ dead cells (Hanayama et al., 2006). As a result, MFG-E8, along with other PS-binding reagents such as the calcium-dependent Annexin V and T-cell membrane protein 4 (TIM-4), has been widely used to study cell death, apoptosis, and EVs (Crowley et al., 2016; Dasgupta et al., 2006).

We have recently developed an *in vivo* labeling technique using MFG-E8 to track and distinguish PS^+^ apoptotic cells from those associated with PS^+^ EVs through imaging flow cytometry (IFC) (Kranich et al., 2020). Our findings indicate that the interaction between PS^+^ EVs and live cells can serve as a powerful marker of disease severity (Rausch et al., 2021) and can modulate immune cell functions (Rausch et al., 2023).

In the present study, we compared various PS-binding reagents to assess their ability to detect free blood EVs, aiming to determine whether differing PS-binding properties could account for the discrepancies reported in the literature. The reagents evaluated included recombinant full-length mouse MFG-E8-eGFP, streptavidin (SA)-tetramerized MFG-E8 C1 domain (C1-Tetramer), and Annexin V, which is widely used in research.

Using imaging flow cytometry (IFC), bead-assisted flow cytometry, and super-resolution microscopy, we observed a predominance of PS^+^ EVs in both murine and human blood. Notably, we found significant differences in the affinities of the PS-binding proteins, which may explain the inconsistencies found in previous studies. We then utilized the most sensitive reagent, C1-Tetramer, to investigate the *in vivo* kinetics and turnover of endogenous EVs, both circulating and cell-bound.

## MATERIALS AND METHODS

### Mice

All mice were housed and bred under specific pathogen-free conditions at the Core Facility Animal Models of the Biomedical Center of the Ludwig-Maximilians-University, Munich. All protocols were approved by the Government of Oberbayern. Female Age-matched mice were used at 8 to 12 week of age. C57BL/6NRj mice were purchased from Janvier (strain C57BL/6). For endogenous EV-labeling mice were injected with 50 ug C1-Tetramer.

### Antibodies/Reagents

See Suppl. Table S3.

### MFG-E8 based PS staining reagents

The C1-Tetramer was produced by mixing biotinylated mC1 (patent publication No. US 2024/0125806 A1) with SA-AF647 (BioLegend), SA-BV421 (BioLegend), SA-AF488 (Biolegend) or SA-CF568 (Biotium) in a 1:5 ratio. After tetramerization, the samples were centrifuged for 30 min at 20,000 g 4°C to remove aggregates. Recombinant MFG-E8-eGFP was produced as previously described (Kranich et al., 2020).

### Blood EV isolation

Mouse blood was collected via heart puncture and coagulation was prevented by addition of heparin (Ratiopharm). Plasma was separated by centrifugation at 1,500 g for 10 min at 4°C. Plasma was diluted to 800 μl in DPBS containing 1x protease inhibitor (Roche) and subsequently centrifuged two more times (2,500 g, 10’, at 4°C and 10,000 g, 10’, at RT) before application to qEV 35-nm columns (Izon Science). Flow-through was collected in 500 μl fractions. EV-containing fractions were determined by NTA, BCA and Western blotting, pooled and concentrated to 300 μl. Human plasma was collected from 5 healthy individuals (3 males, 2 females – age 21-60) into S-Monovette® Natrium Heparin tubes and processed as described above for murine samples. The study protocol (Project-Nr. 18-0415) was approved by the Institutional Review Board of the Medical Faculty of LMU Munich.

### BCA, SDS-PAGE and Western blotting

Protein contents were measured using a BCA protein assay kit (Thermo Scientific Pierse, Rockford, IL, USA) according to manufacturer’s instructions, using BSA as a standard. For Western blotting, the 12 EV fractions were pooled in pairs and concentrated to 200 μl. 20 μl of each sample were denatured at 95°C for 5 min in the presence of 1x loading buffer (50 mM Tris-HCL pH 6.8, 100 mM DTT, 2% SDS, 0.1% Bromophenol blue, 10% glycerol). Samples were loaded onto 12% SDS-PAGE for protein separation. Proteins were transferred onto nitrocellulose blotting membrane (Amersham, Cytiva) in a Mini Trans-blot ® cell system (Bio-Rad). After transfer, membranes were blocked with 1x Casein blocking buffer (Sigma-Aldrich) for 90 min at RT. Primary antibodies were diluted in 1x Casein blocking buffer and incubated overnight at 4°C. After washing with TBS-T, secondary antibodies were diluted in 1x Casein blocking buffer were added for 90 min at RT. Following washing with TBS-T, Western Lightning® Plus-ECL (PerkinElmer) reagents were used for detection of protein bands in an iBright^TM^ 1500 imager (ThermoFisher).

### C1-Tetramer EV labeling for Transmission electron microscopy

The C1-Tetramer-gold for TEM imaging was produced by mixing biotinylated mC1 with SA conjugated with 6 nm gold particles (Aurion) in a 1:5 ratio. Isolated murine plasma EVs (500 µl) were labelled with C1-Tetramer-gold or streptavidin-gold for 90 min at RT at a final concentration of 7 ng/µl. Afterwards, EVs were fixed with 2.5% glutaraldehyde (EM-grade, Sciences Services) for 10 min at RT and the EVs were diluted with 20 ml 1x TBS followed by concentration of the sample to the starting volume of 500 µl using Vivaspin® 30 MWCO tubes (Sartorius). Prior to staining, 200-mesh TEM grids with carbon-coated Formvar film were rendered hydrophilic by glow discharging for 30 seconds. Next, 1.5 µl of the purified EVs were transferred onto the grids and incubated for 2 minutes to allow adsorption to the grid surface. Excess liquid was blotted with filter paper, and the grids were washed by adding a drop of distilled water for 30 seconds, followed by blotting the excess water. The samples were then stained in a drop of 1% (w/v) aqueous uranyl acetate for 30 seconds in the dark. Excess uranyl acetate solution was blotted, and the grids were air-dried. Samples were imaged using a JEM-1400 Plus (JEOL) transmission electron microscope operated at an accelerating voltage of 120 kV, equipped with a TemCam XF416 digital camera (TVIPS).

### Superresolution microscopy dSTORM

For direct stochastic optical reconstitution microscopy (dSTORM) analysis, 15 μl of EVs at 1-5×10^10^ particles/ml were stained overnight at 4°C with antibodies and the C1-Tetramer-AF647 in the presence of 0.5% bovine serum albumin and Fc receptor blocking diluted in filtered DPBS. Purified antibodies were labelled in-house with dSTORM compatible dyes using Mix-n-Stain™ CF™488 and CF™568 kits according to the manufacturer’s instructions. Two different negative controls were included in all experiments. Firstly, EVs stained with isotype control antibodies and SA-AF647. Secondly, antibodies diluted in DPBS without EVs. The next day, flow chambers were assembled consisting of SA-coated glass slides (Ibidi) and 1.5H cover slides. Flow chambers were either coated with 0.3 mg/ml with commercially available C1-Tetramer (Apo-Monomer, Biolegend #427405) for PS-based capture or with CD9/CD81-biotin at a final concentration of 0.25 mg/ml, both for 15 mins at RT. After coating, flow chambers were washed twice with 100 μl filtered DPBS before proceeding to EV capture for 45 min at RT. Washing was repeated as previously mentioned and afterwards EVs were fixed with 4% paraformaldehyde for 10 mins at RT, followed by another washing step. Freshly prepared BCubed STORM-imaging buffer (ONI, Oxford Nanoimaging) was added prior to image acquisition on a temperature-controlled Nanoimager S Mark II microscope from ONI. Images were taken in dSTORM mode acquired sequentially using the total reflection fluorescence (TIRF) illumination (calculated evanescent field penetration depth was >200 nm). Before imaging, channel mapping was calibrated using 0.1 µm TetraSpeck beads (T7279, Thermo Fisher Scientific). Superresolution images were filtered using the NimOS software (v.1.18.3, ONI) and data have been further processed with the Collaborative Discovery (CODI) online analysis platform from ONI. Post-processing data analysis included correction for drift, temporal grouping and filtering to remove suboptimal localisation and improve overall localisation precision. For cluster analysis HDBSCAN was used and at least 15 localisations were required to constitute a cluster.

### Imaging flow cytometry for EV analysis

The imaging flow-cytometer (IFC) ImageStream^TM^ MKII (Cytek) requires small volumes of EV sample (25-200 µl). We opted for a staining volume of 60 µl and a final volume of 120 µl, adjusting buffer/reagent/antibody concentrations appropriately, since no washing steps can be performed following the labeling. The primary purpose of dilution is to balance between timely acquisition and potential complications associated with too high concentration - for instance “swarm formation.”

All EV or liposome preparations analyzed by IFC were stained either with CFSE (carboxyfluorescein succiminidyl ester, eBiosciences) or CellTrace^TM^ violet (Thermo Fisher) which are commonly used for labeling of membranous vesicles such as EVs. Liposomes with defined size and DOPS:DOPC content were purchased from Encapsula NanoSciences. Prior to staining, EV or liposome concentration was determined using NTA analysis and adjusted to 1-5×10^10^ particles/ml. All staining reagents were centrifuged at 18,000g for 30 min at 4°C. 1 µl of CFSE or CTV were added to 40 µl of sample material to reach a final concentration of 10 µM. After addition of the membrane dyes, samples were incubated for 1 h at RT protected from light. Antibodies or staining reagents were added to the samples (for concentrations see Supplementary Table S1 and S2) in a total volume of 60 µl (to reach 60 µl filtered PBS was added if necessary). For Annexin V staining, 6 µl of Annexin binding buffer (Annexin V 10x binding buffer, BD Pharmingen) were added to the samples while the total volume of 60 µl remained the same. Subsequently, samples were incubated for 1 h at RT, protected from light. Afterwards, samples were split in half and 2.5% (v/v) NP-40 was added to the detergent controls whereas the same volume of PBS was added to the samples. Prior to IFC measurements, samples were diluted to 120 µl with filtered PBS or 120 µl filtered PBS containing 1x Annexin V binding buffer. Aggregate controls contained the same final concentration of antibodies, staining reagents or membrane dyes diluted in filtered PBS. Antibody specificity was tested for all mABs using appropriate isotype controls or SA-fluor for the C1-Tetramer. As negative controls, fluorescence minus one, unstained EVs, reagent/buffer and detergent controls were used. For all panels used here, serial dilutions were performed to exclude bias stemming from “swarm detection”. According to the MISEV guidelines (Welsh et al., 2024) comprehensive list of all staining reagents, concentrations and methodological procedures can be found in the supplement (Table S1 and S2).

Prior to single vesicle analysis, we tested the ability of our ImageStream^TM^ MKII to detect single-size populations of fluorescent sub-micron beads by measuring two commercially available mixtures of FITC-fluorescent polystyrene (PS) beads of known sizes (Megamix-Plus FSC – 900, 500, 300 and 100 nm, and Megamix-Plus SSC – 500, 240, 200, 160 nm). Next, we mixed both bead sets in a 1:1 ratio (‘Gigamix’) and performed acquisition and tested the ability of the ImageStream^TM^ MKII to discern all seven fluorescent bead populations, as well as the 1 µm-sized Speed Beads (SB), via the FITC (Ch02) and side scatter (SSC - Ch06) intensities. For subsequent EV analysis, we made use of CFSE or CTV membrane labeling and gated on low scatter CFSE/CTV^+^ events. Acquisition of EVs was performed with fluidics set at low speed, sensitivity set to high, magnification at 60x, core size 7 µm, and the “Remove Beads” option unchecked prior to every acquisition. The ImageStream^TM^ MKII was equipped with the following lasers run at maximal power to ensure maximal sensitivity: 405 nm (120 mW), 488 nm (200 mW), 561 nm (200 mW), and 642 nm (150 mW). Upon each startup, the instrument calibration tool ASSIST® was performed to optimize performance and consistency. Two channels (Ch01 and Ch09) were set to bright-field (BF), permitting spatial coordination between cameras.

### Preparation of single cells suspensions for flow cytometry

Single-cell suspensions of the spleen were prepared by meshing the organs through a nylon mesh followed by erythrocyte lysis using an ammonium-chloride-potassium buffer. 5 × 10^6^ cells were blocked with Fc block (10 min on ice) and stained for imaging flow cytometry with appropriate antibody mixes (25 min on ice), after the staining procedure cells were washed twice with DPBS and finally resuspended in 50 μl for IFC or 500 μl for analysis on the BD FACSCanto II (BD Biosciences). When staining with Annexin V, cells were stained, washed and resuspended in 1x Annexin V buffer (BD Pharmingen) diluted with PBS. For IFC acquisition, cells were analyzed at low speed and 60x magnification on an ImageStream^TM^ MKII. PS^+^ cells were identified using FMO controls and for IFC analysis apoptotic and EV^+^ cells were digitally sorted using a machine-learning based convolutional autoencoder (CAE) as previously described^1^.

### Nanoparticle tracking analysis

Particle number and size distribution of EVs were determined by NTA using the ZetaView PMX110 instrument (ParticleMetrix). Eleven positions were measured with three reading cycles with temperature control at 21°C enabled. Preacquisition parameters were sensitivity = 75, shutter speed = 50, frame rate = 30 fps, trace length = 15. Postacquisition parameters were minimum brightness = 20, pixels size = 5 to 1,000.

### Multiplex bead-based EV surface protein profiling

Human or murine purified EV fractions were analyzed using MACSplex exosome kit (MACSplex Exosome kit human/mouse, Miltenyi) for multiplex analysis by flow cytometry. All sample were adjusted to 1-2×10^10^ particles per assay and incubated with MACSplex exosome capture beads for 18 h on an orbital shaker at 450 rpm at room temperature.

Beads were washed according to the manufacturer’s protocol with either PBS/BSA 1%(w/v) or for Annexin V staining PBS/BSA 1%(w/v)/ 1x Annexin V binding buffer. For staining of the captured EVs, different concentrations of Annexin V-AF647 (Biolegend) or the C1-Tetramer-AF647 were used. As controls served beads without EVs, beads with EVs but no PS-labeling (PS FMO) or beads with EVs stained with SA-AF647. All staining reagents were incubated with the samples for 1 h on an orbital shaker at 450 rpm and RT. Afterwards, samples were washed twice and finally resuspended in 300 μl PBS/BSA 1%(w/v) (with or without Annexin V binding buffer). All samples were acquired on a BD FACSCanto II (BD Biosciences). Geometric mean fluorescent intensities (MFIs) for all capture bead subsets were background-corrected by subtracting respective MFI values from matched non-EV containing buffer controls. MFI values below those obtained for EVs stained with SA-AF67 of each respective sample are reported as ‘not detected’ for C1-Tetramer staining. For Annexin V, an Annexin V FMO control was used for the same purpose.

### Statistics

For statistical analysis, the PRISM software (GraphPad Software) was used. Significance was analyzed using one-way ANOVA test unless stated otherwise, ns P>0.05. Graphs show average ±SEM. Dissociation constants (kD) were calculated by fitting non-linear regression curves with Hill slope equation in PRISM, goodness of the fit was evaluated by R^2^ values which ranged from 0.84 to 0.99. The half-life of *in vivo* labelled plasma EVs was determined by fitting a non-linear one phase decay curve in PRISM, goodness of the fit was evaluated by R^2^ (0.91).

## RESULTS

### Comparison of PS-binding of different PS-specific reagents

We have developed recombinant PS-binding proteins, including (i) full-length MFG-E8, (ii) MFG-E8 C1C2, which lacks the EGF and PT domains, and (iii) multimerized MFG-E8 C1 domains (C1-Tetramer; Fig. 1A), to differentiate between PS^+^ live cells associated with PS^+^ extracellular vesicles (EVs) and authentic PS^+^ apoptotic cells (Kranich et al., 2020; Rausch et al., 2023; Rausch et al., 2021). Historically, PS-binding proteins like Annexin V and purified bovine MFG-E8 have been applied to label EV preparations, though these often produced heterogeneous results. Considering the variability in PS content (Groß et al., 2024) and EV sizes, we aimed to assess whether different reagents are equally effective for detecting PS^+^ EVs.

**Fig. 1.**
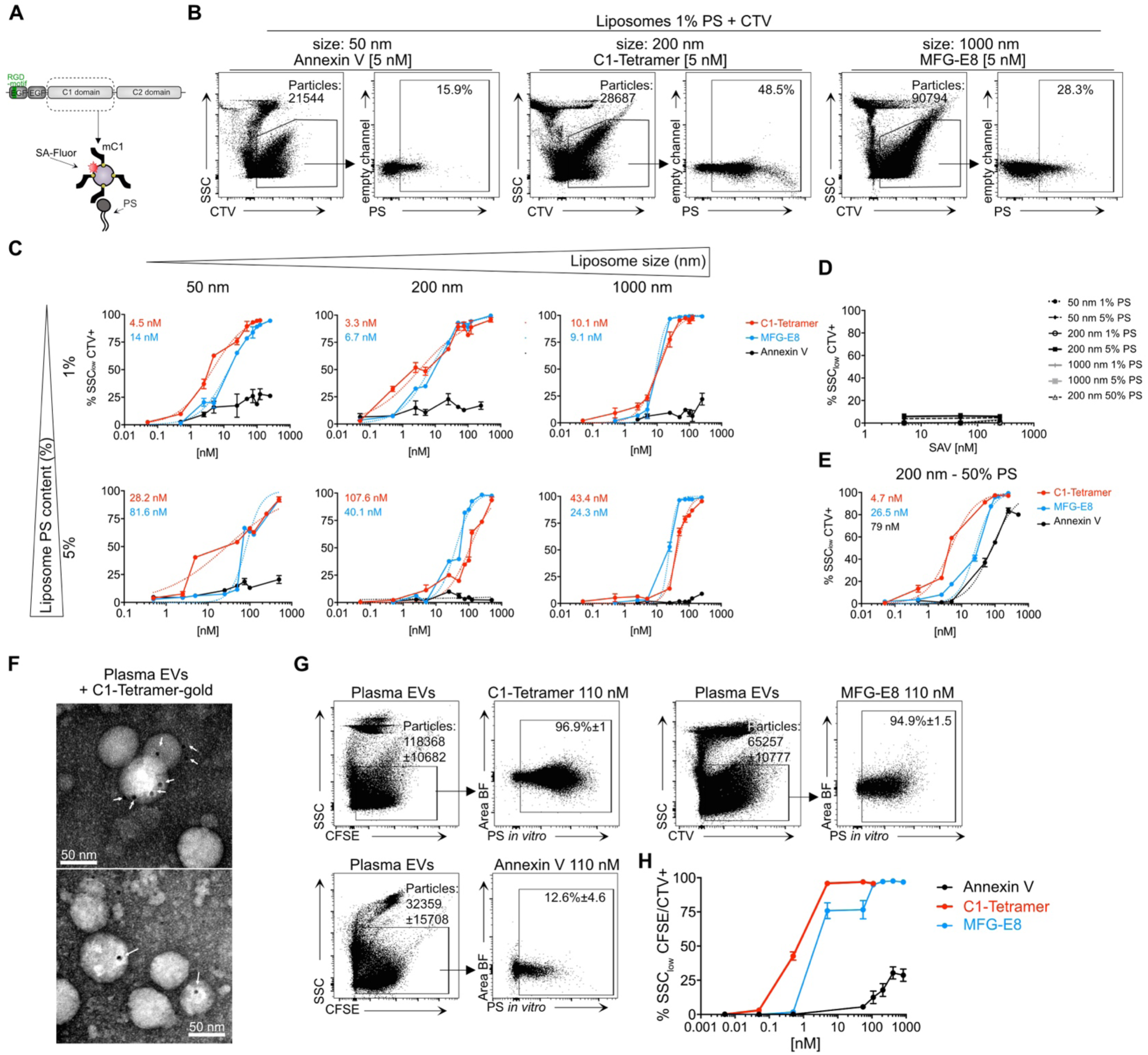
MFG-E8-based PS-binding reagents efficiently stain small liposomes with low PS content and murine plasma EVs. **(A)** Schematic representation of full-length MFG-E8 domains and the C1-Tetramer produced by tetramerization of the monomeric biotinylated C1 domain (mC1) with fluorescently conjugated streptavidin (SA-Fluor). **(B)** Imagestream^TM^ analysis of liposomes (Encapsula), shown is the exemplary gating strategy for 50, 200 and 1000 nm sized liposomes, using the membrane dye cell-trace violet (CTV) and excluding high scatter events. (**C)** Liposomes of defined size and lipid ratio (DOPC:DOPS 99:1 or 95:5), were used to determine the affinity of Annexin V-AF488 (black), full-length MFG-E8-eGFP (blue) or the C1-Tetramer-AF488 (red). All samples were measured as technical duplicates or triplicates. **(D)**, Binding specificity of the C1-Tetramer was assessed by incubating liposomes with increasing concentrations of SA-AF488. **(E)**, Liposomes (200 nm) with high PS content (DOPC:DOPS, 1:1) labelled with distinct concentrations of Annexin V-AF488 (black), full length MFG-E8-eGFP (blue) or the C1-Tetramer-AF488 (red). Dissociation constants (kD) were calculated by fitting non-linear regression curves with Hill slope equation. (**F)** Transmission electron microscopy images of C1-Tetramer-gold labelled murine plasma EVs. White arrows indicate gold particles. (**G),** Sensitivity of different PS-binding proteins for PS+ murine plasma EVs was examined by IFC, exemplary gating on EVs is shown using the membrane dye carboxyfluorescein succinimidyl ester (CFSE) or CTV and excluding high scatter events. **(H)**, PS-labelling efficiency of membrane particles at distinct concentrations of Annexin V (AF647), C1-Tetramer (AF647) or MFG-E8-eGFP (n=3).

To this end, we first conducted experiments using liposomes of various sizes (50, 200, and 1000 nm) with defined PS contents of 1% and 5% (Fig. 1B, C). We then analyzed the labeling efficiency of these liposomes by fluorescently conjugated PS-binding proteins using imaging flow cytometry, following an initial calibration (Suppl. Fig. 1A), as previously described (Woud et al., 2022). For all single particle flow cytometry experiments, we adhered to MiFlowCyt recommendations for controls, with detailed tables (Suppl. Table 1 and 2) and figures available in the supplement (Welsh et al., 2024).

For liposome labeling, we used the membrane dye CellTrace Violet (CTV) to determine the percentage of PS positivity across all detected particles (Fig. 1B and Suppl. Fig. 1B-F). This analysis revealed notable differences between MFG-E8 derivatives and Annexin V; the latter failed to label significant portions of any liposome populations tested (Fig. 1C). Furthermore, Annexin V’s labeling efficiency plateaued at 25% for small liposome types, even at higher concentrations. In contrast, both full-length MFG-E8 and the C1-Tetramer achieved nearly 100% labeling efficiency, regardless of liposome size or PS content (Fig. 1C). Notably, the C1-Tetramer demonstrated higher sensitivity (kD) than full-length MFG-E8 for the smallest (50 nm, 1% and 5% PS) and medium-sized (200 nm, 1% PS) liposomes. This finding aligns with prior research indicating that liposome curvature and PS content influence full-length MFG-E8 binding (Carman et al., 2023; Shi et al., 2004). Importantly, the C1-multimer, comprising the minimal PS-binding C1 domain of MFG-E8, extends applicability to even the smallest vesicles with low PS content (Fig. 1C). Given the failure of Annexin V to bind to liposomes with low PS frequency, we next included 200 nm liposomes with 50% PS content in our analysis (Fig. 1E). Indeed, Annexin V could label close to 90% of the high PS frequency liposomes, albeit still with the lowest affinity compared to MFG-E8 and the C1-Tetramer.

We then tested these PS-binding proteins on EVs purified from murine and human blood, where PS levels are typically very low (Groß et al., 2024), making reagent sensitivity particularly crucial. Plasma EVs were isolated from mouse blood through sequential centrifugation and size-exclusion chromatography (SEC), with EV-containing fractions identified by nanoparticle tracking analysis (NTA), protein quantification (bicinchoninic acid assay), and Western blotting (Suppl. Fig. 1G,H). The isolated plasma EVs had a mean size of 152 ± 10.2 nm, which remained unchanged when EVs were labeled with the C1-Tetramer (146 ± 5.6 nm; Suppl. Fig. 1I). To validate the binding specificity of the C1-Tetramer, SA-gold was used to generate C1-tetramers for transmission electron microscopy (TEM), which confirmed the presence of PS^+^ EVs, showing round and cup-shaped vesicular structures (Fig. 1F). TEM was also employed using a negative control stain with SA conjugated to gold particles (Suppl. Fig. 1J).

We then analyzed EVs by imaging flow cytometry (IFC) (Welsh et al., 2024). To evaluate surface PS positivity by IFC, EVs were defined as membrane dye-positive (CFSE or CTV) and side scatter (SSC) low (Fig. 1G), with detergent sensitivity further confirming their membranenous nature (Suppl. Fig. 2 B-D). EVs were subsequently stained with increasing concentrations of Annexin V, recombinant full-length murine MFG-E8, or the C1-Tetramer to assess the frequency of PS^+^ EVs (Fig. 1H).

Our findings showed significant differences in PS^+^ EV detection across reagents, consistent with prior liposome results. At the highest concentration, Annexin V labeled only about 28% of EVs, while both the C1-Tetramer and MFG-E8 detected nearly all EVs (approximately 95-97%, Fig. 1H). Notably, the C1-Tetramer achieved maximum detection levels at concentrations 166 times lower than Annexin V, whereas MFG-E8 reached similar detection levels at concentrations 22 times higher than the C1-Tetramer, indicating the latter’s superior affinity (Fig. 1H).

Previous studies, including our own, have demonstrated that MFG-E8 derivatives and Annexin V exhibit similar detection capacities for PS^+^ dead cells (Hanayama et al., 2002; Kranich et al., 2020; Rausch et al., 2023) (Suppl. Fig. 2E). This contrasts with our findings above, where Annexin V showed reduced detection of PS^+^ EVs but it could be explained by the dependence of Annexin V binding on PS frequency as demonstrated by using liposomes (Fig. 1E). To further investigate, we used heat-killed splenocytes to induce cell death and promote PS externalization. The cells were then stained with varying concentrations of PS-binding reagents. When gating on all PS^+^ splenocytes (both Live-Dead (LD)^+^ and LD^−^), the C1-Tetramer stained a greater proportion of cells (81.3 ± 2.2%) compared to Annexin V (66.9 ± 2.3%) at the highest concentration (Suppl. Fig. 4A-C). Interestingly, MFG-E8 stained only 45 ± 1.3% of splenocytes, less than Annexin V.

However, when focusing on necrotic (LD^+^) cells, differences among Annexin V, MFG-E8, and the C1-Tetramer were minimal, with detection rates of 99.5 ± 0.1%, 97.9 ± 0.1%, and 98.9 ± 0.1% at the highest concentration, respectively. This suggests that, while reagent efficacy varies across PS^+^ cell populations, it remains comparable for necrotic cells.

This observation also included a higher median fluorescence intensity (MFI) for PS on LD^+^ cells compared to LD^−^ cells across all staining reagents (Suppl. Fig. 4C). In contrast, differences in PS-binding protein sensitivity became apparent when focusing on LD^−^ splenocytes. Among these reagents, the C1-Tetramer showed the highest sensitivity, followed by Annexin V and MFG-E8. LD^−^ splenocytes encompass apoptotic and EV-decorated cells, which generally exhibit less PS exposure than necrotic cells (Suppl. Fig. 4C) (Kranich et al., 2020).

Together, these findings highlight the confounding effects of using PS-staining reagents with varying sensitivities, a factor especially pronounced in contexts with low PS expression, such as on blood EVs. For Annexin V, only a subset of EVs was detected, likely representing EVs with high PS exposure, while MFG-E8 appears more effective in binding EVs with both high and low PS frequencies (Fig. 1C, E and H). Interestingly, PS exposure emerges as a dominant feature of plasma EVs when using the C1-Tetramer and, despite its lower sensitivity, also with MFG-E8.

### PS exposure is a common characteristic across all EV populations in mice

Given the heterogeneous nature of blood-derived EVs, we aimed to further characterize PS^+^ EVs and differentiate them from potential contaminants. Tetraspanins such as CD9, CD63, and CD81, which are enriched on exosomes and other EVs compared to their host cells, are often considered universal EV markers (Raposo & Stoorvogel, 2013). Therefore, we examined PS positivity among tetraspanin^+^ EV populations using CD9/CD81 antibodies and imaging flow cytometry (IFC) (Fig. 2A). When gating on tetraspanin^+^ EV subsets, the proportion of PS^+^ events ranged from 82.8% to 99.2%, representing most of the distinct EV populations.

**Fig. 2:**
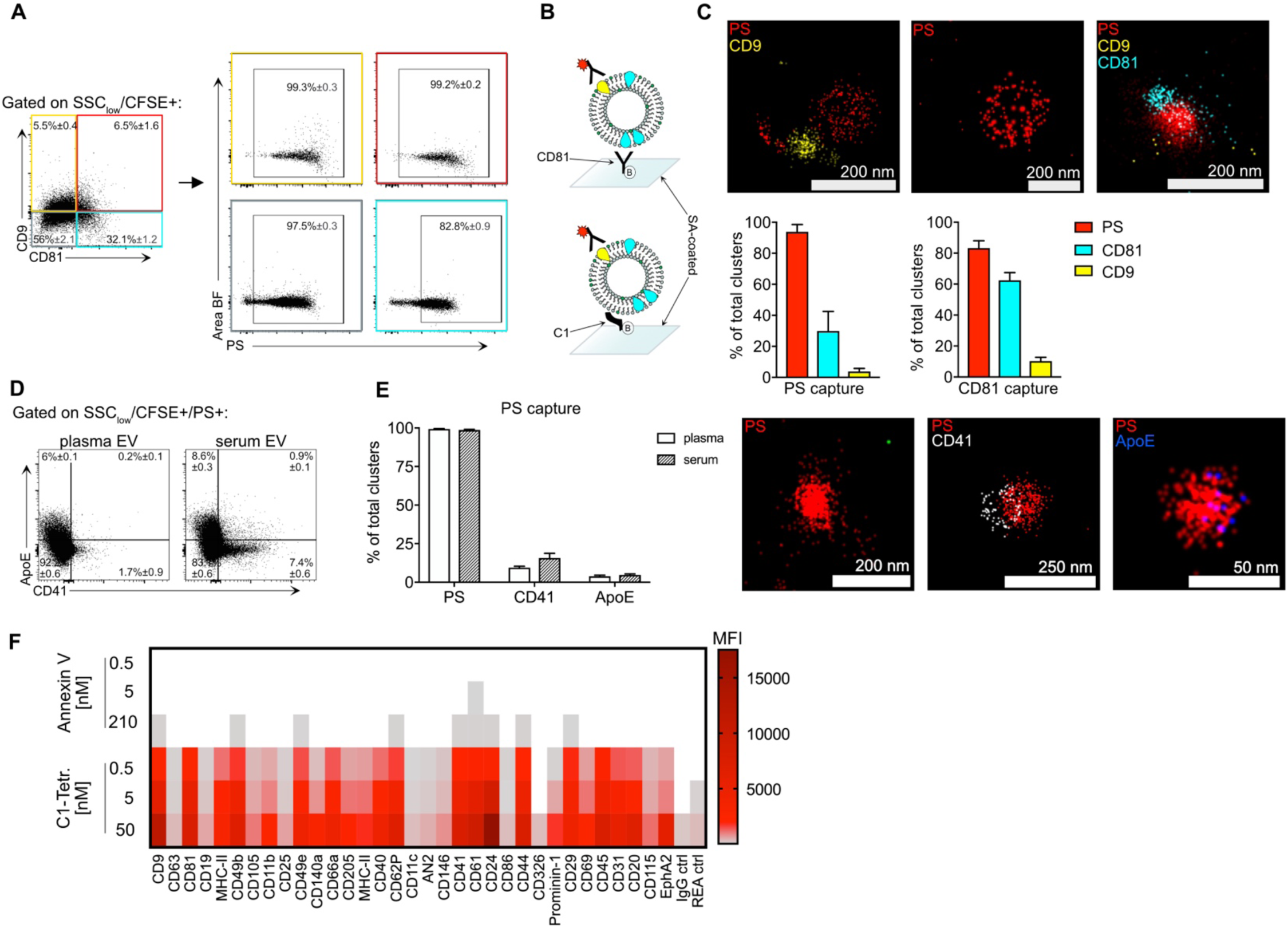
C1 tetramers stain various EV populations. **(A)** IFC anaylsis for tetraspanins CD9 and CD81 on murine plasma EVs (n=3), PS positivity of distinct EV subsets was examined. **(B)** Schematic representation of PS vs CD81 based capturing for superresolution microscopy of EVs, briefly, stained EVs were captured onto SA-coated glass slides using biotinylated mC1 or antibodies. **(C)** Plasma EVs (n=5) stained for tetraspanins CD9/CD81 and PS were examined by superresolution microscopy and cluster analysis using different capturing reagents, exemplary dSTORM micrographs are shown in the upper row. Bar graphs show percentages of PS+ (red), CD81+ (green) and CD9+ (yellow) clusters ±SEM after PS capturing (left) or CD81 capturing (right) **(D)-(E)** IFC analysis of SSC_low_/CFSE+/PS+ murine plasma or serum EVs (n=3) for a platelet marker CD41 and a lipoprotein marker ApoE. Bar graphs show percentages of PS+, CD41+ and ApoE+ clusters in plasma and serum. Error bars represent SEM. **(F)** Surface exposure of PS (rows) analyzed by bead-assisted flow cytometry, using the C1-Tetramer-AF647 or Annexin V-AF647 for PS staining. Distinct EV subpopulations are shown which were captured by bead-coupled antibodies targeting the indicated proteins (columns). Color coding indicates MFI for AF647 of the respective bead population, MFI values were background corrected by a ‘no EV’ blank control and for MFI values obtained for SA-AF647 or an Annexin V FMO for the C1-Tetramer and Annexin V staining, respectively. Displayed are means of biological replicates (n = 3).

To confirm these results, we employed direct stochastic optical reconstruction microscopy (dSTORM), achieving a resolution of ∼20nm, to evaluate PS positivity in specific EV populations. We immobilized plasma EVs on glass slides using either anti-CD81 antibodies or C1-tetramers (Fig. 2B). Cluster analysis, conducted with isotype control antibodies and buffer as negative controls (Suppl. Fig. 5D), revealed that 94±2.4% of EVs captured with C1-Tetramers were PS^+^, including subsets double- or triple-positive for CD81 (30±6.3%) and/or CD9 (4±1%). When capturing EVs with anti-CD81 antibodies, the total proportion of clusters positive for CD81 (64±5.7%) and CD9 (12±2.2%) increased, though not all clusters were tetraspanin^+^. This may be due to tetraspanin-enriched microdomains on EVs, as previously described (Andreu & Yánez-Mó, 2014), potentially causing limited epitope availability or steric hindrance during EV capture. Like PS capture, nearly all EVs captured with anti-CD81 were also PS^+^ (80±3.8%), corroborating IFC findings.

To assess possible contamination of lipoprotein particles, such as chylomicrons, we used IFC and super-resolution microscopy to quantify ApoE, a marker found on murine HDL, LDL, VLDL, and chylomicrons (Gordon et al., 2015). We also analyzed EVs carrying the platelet marker CD41 to determine whether blood preparation methods (serum vs. plasma) could promote artificial release of platelet EVs. We observed an increased frequency of CD41^+^ PS^+^ events in serum EV preparations (Fig. 2D and Suppl. Fig. 5E-G) following coagulation, whereas in plasma EVs, only a small fraction of PS^+^ events were CD41^+^ (1.7±0.9%) or ApoE^+^ (6±0.1%). Super-resolution microscopy supported these observations, showing ApoE present in only 4±0.6% and 5±0.6% of clusters in plasma and serum EV preparations, respectively (Fig. 2E).

To further investigate PS labeling differences, we compared Annexin V and C1-Tetramer using a multiplex bead assay that analyzes 37 distinct EV populations based on surface protein capture. Exposed PS on captured plasma EVs was detected with Annexin V-AF647 or C1-Tetramer-AF647 (Fig. 2F). Staining with C1-Tetramer revealed PS signal above background for all EV populations except CD326^+^ EVs, regardless of concentration. This supports the notion that PS exposure is a predominant feature of diverse EV types.

However, signal intensities among populations may reflect differences in vesicle capture rather than PS content alone. In contrast, PS staining with Annexin V required a concentration four times higher than C1-Tetramer, and even then, only one-third of EV bead populations showed detectable PS. Notably, the Annexin V-positive populations aligned with those showing the highest C1-Tetramer signal intensities, such as CD9.

Overall, our results demonstrate that PS exposure is a defining feature of blood EV populations. Importantly, in our isolated plasma EV preparations, only a small fraction (4– 6%) of PS^+^ particles could be attributed to common contaminants like lipoproteins or platelets. Additionally, this study highlights the labeling bias introduced by varying PS-binding protein affinities, as evidenced by differing efficiencies of Annexin V and C1-Tetramer in EV population multiplex assays.

### The majority of human plasma EVs similarly exhibit PS exposure

To determine whether human plasma EVs exhibit similar PS-positivity as observed in mice, we analyzed EVs isolated from human plasma. EV isolation and validation followed the protocol established for murine samples (Supplementary Fig. 7A, B), yielding EVs with a mean size of 131±11.8 nm as determined by nanoparticle tracking analysis (NTA) (Suppl. Fig. 7C).

Consistent with murine data, the majority of SSC_low_CTV^+^ events in human plasma EVs were PS^+^ (84.7±1.6%). Upon comparing CD9^+^ and CD9^−^ EV populations by IFC, we observed slightly lower PS levels in CD9^+^ EVs (71.9±1.9%) than in CD9^−^ EVs (89.6±0.9%; Fig. 3A and Suppl. Fig. 7D-F). This trend was further confirmed using super-resolution microscopy, where 84±4.2% of the CD9-captured EV clusters were PS-positive (Fig. 3B).

**Fig. 3:**
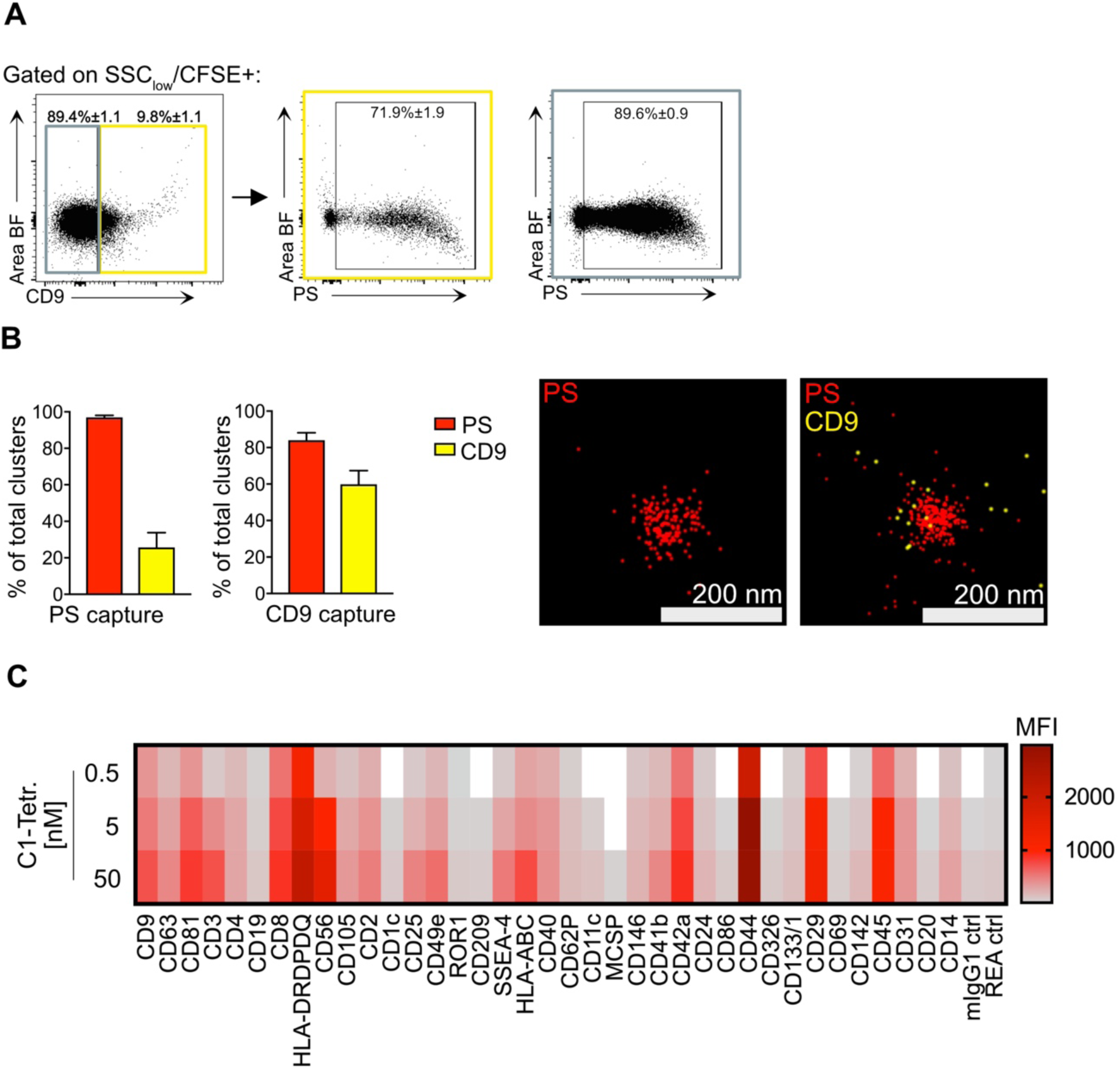
Almost all human plasma EVs are PS^+^. **(A)** Human plasma EVs (n=5) were isolated and analysed by IFC for PS positivity of CD9+ and CD9-EVs. SSC_low_CFSE+CD9+ and CD9-EVs were gated (left) followed by gating of PS+ CD9+ and CD9-EVs. Percentages ±SEM of PS+ EVs are depicted within the gate. **(B)** Superresolution microscopy of human EVs (n=3) stained for the tetraspanin CD9 and PS. Bar graphs show percentages of PS+ (red) and CD9+ (yellow) clusters ±SEM Exemplary dSTORM micrographs are shown to the right. **(C)** Surface exposure of PS (rows) analysed by bead-assisted flow cytometry, using the C1-Tetramer-AF647 for PS staining. Distinct EV subpopulations are shown which were captured by bead-coupled antibodies targeting the indicated proteins (columns). Fluorescently conjugated streptavidin (SA) served as a negative control to test specificity of C1-Tetramer labeling. Colour coding indicates MFI for AF647 of the respective bead population, MFI values were background corrected by a ‘no EV’ blank control and for MFI values obtained for SA-AF647. Displayed are means of biological replicates (n = 3) for the C1-Tetramer staining.

A multiplex analysis of human plasma EVs detected PS signals above background in 36 out of 37 EV bead populations, except for EVs captured using anti-melanoma-associated chondroitin sulfate proteoglycan (MCSP; Fig. 3C and Suppl. Fig. 8). This multiplex assay included general EV markers, such as the tetraspanins CD9, CD63, and CD81, as well as markers indicative of cellular origin (e.g., CD3, CD4, CD8, CD11c, CD45) and cell activation (e.g., CD44). These findings highlight both the heterogeneity and the prevalence of PS^+^ EVs in human plasma samples, mirroring observations in murine models.

### Kinetics of Endogenous PS^+^ EVs Determined by In vivo Labelling

Based on our findings, we concluded that PS is widely exposed on nearly all plasma EVs in both mice and humans. We sought to utilize this characteristic to study the turnover of endogenous EVs *in vivo* using the C1-Tetramer. In previous studies, we demonstrated the effectiveness of both MFG-E8 and the C1-Tetramer for *in vivo* labeling of cell-bound EVs, and here we extend their application to analyze circulating unbound EVs simultaneously (Fig. 4A).

**Fig. 4:**
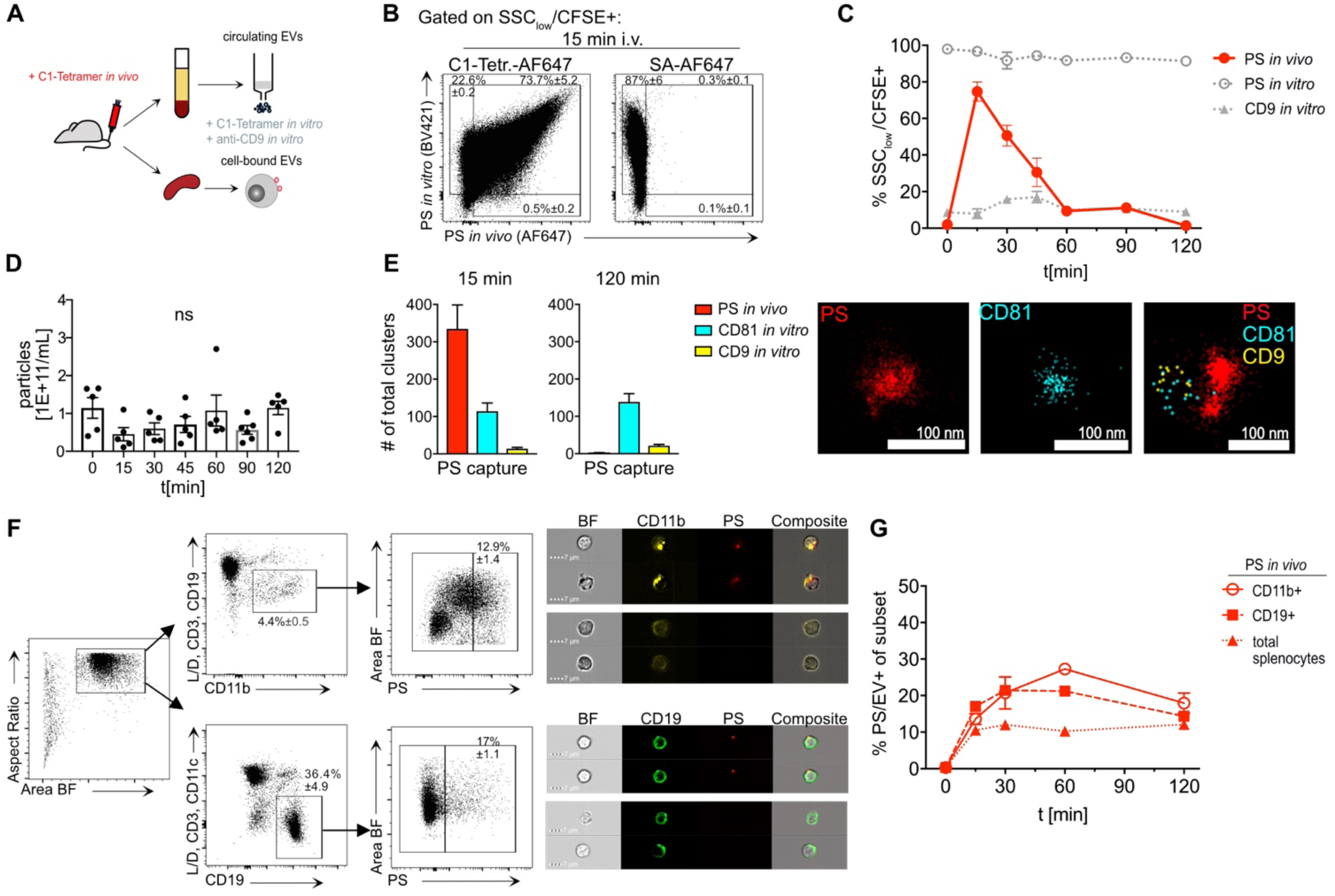
In vivo measurement of EV turnover. **(A)** Schematic illustration of *in vivo* PS-labeling. 50 μg of the C1-Tetramer were i.v. injected and plasma EVs as well as splenocytes were collected at different timepoints and analyzed by IFC. **(B)** Dot plots show *in vivo* vs. *in vitro* PS-labeled plasma EVs after 15 min post-i.v. injection of the C1-Tetramer-AF647 (left) or SA-AF647 as a negative control (right). **(C)** Kinetic of *in vivo* PS-labeled EVs analyzed by IFC at 15, 30, 45, 60, and 120 min after C1-tetramer injection. Graphs shows percentage of *in vivo* (red) and in vitro (grey, open circles) PS-stained and in vitro CD9-stained (grey, closed triangles) ±SEM SSC_low_CFSE^+^ EVs at different timepoints. **(D)** Nanoparticle tracking analysis quantification of particle concentrations in plasma samples at different timepoints post-injection of the C1-Tetramer. **(E)** *In vivo* PS-labeling of murine plasma EVs at two different timepoints analysed by superresolution microscopy. Bar graphs show number of *in vivo* stained PS^+^ (red), in vitro stained CD81^+^ (cyan) and CD9^+^ (yellow) EV clusters ±SEM. Exemplary dSTORM micrographs are shown to the right. **(F)** IFC analysis of splenocytes, shown is the gating strategy for splenic PS^+^CD19^+^ B cells and PS^+^CD11b^+^ monocytes/macrophages, exemplary images for EV/PS^+^ cells are shown to the right. **(G)** Graph shows kinetic of *in vivo* labeled PS/EV^+^ splenocyte populations at 0, 15, 30, 60 and 120 min as measured by IFC.

First, we examined whether C1-Tetramer-labeled EVs could be detected using imaging flow cytometry (IFC) following intravenous (i.v.) injection and subsequent EV isolation. To avoid potential bias from residual free C1-Tetramer, we tested the stability of the labeling (Supplementary Fig. 9G-I). To verify that free C1-Tetramer was fully removed after size exclusion chromatography (SEC), we pre-labeled two separate EV samples with C1-Tetramer conjugated to distinct fluorophores prior to SEC isolation. After a 90-minute co-incubation of the individually labeled EVs, no co-staining was observed, confirming the effective removal of free C1-Tetramer.

Next, we injected mice with 50 μg C1-Tetramer and isolated plasma EVs 15 minutes later. We observed that 74±5.2% of detected EVs were labelled. However, in vitro PS counterstaining revealed that approximately 22.6±0.2% of the EVs were unlabeled *in vivo* (Fig. 4B). This discrepancy may be due to either reduced labeling efficiency *in vivo*, potentially influenced by dynamic blood flow, or rapid clearance. Nevertheless, the majority of EVs were labeled, demonstrating the suitability of C1-Tetramer injection for studying blood EV turnover.

Using this approach, we assessed the half-life of PS^+^ plasma EVs via IFC at various time points. *In vivo*-labeled PS^+^ EVs were completely cleared from circulation within 2 hours, with a calculated half-life of 29 min (Fig. 4C). Importantly, *in vivo* PS-labeling did not affect the total population of PS^+^ plasma EVs, as shown by *in vitro* counterstaining for PS and CD9, which indicated stable EV composition over time. Similarly, nanoparticle tracking analysis (NTA) at all time points revealed no changes in EV abundance (Fig. 4D). We further confirmed *in vivo* labeling at 15 and 120 minutes using super-resolution microscopy on PS-captured EVs (Fig. 4E, Suppl. Fig. 9F). At 15 minutes, most EV clusters were PS-labeled *in vivo*, whereas by 120 minutes, PS labeling was undetectable. Notably, CD81^+^ and CD9^+^ EV cluster counts remained constant over time.

Finally, we compared the turnover of circulating plasma EVs with that of cell-bound EVs isolated from the spleen (Fig. 4F, G). After i.v. injection of the C1-Tetramer or streptavidin-AF647 as a negative control (Suppl.Fig. 10B), we isolated splenocytes for IFC analysis. PS^+^ cells were predominantly EV^+^ with fewer than 1% apoptotic cells, as determined by a convolutional autoencoder machine-learning model (Supplementary Fig. 10A) (Kranich et al., 2020).

We focused on splenic B cells (CD19^+^) and monocytes/macrophages (CD11c^+^) due to their high PS/EV positivity (Kranich et al., 2020). Interestingly, we found that the turnover of cell-bound EVs in spleen cells was prolonged compared to circulating EVs. PS/EV^+^ splenocyte frequency remained virtually unchanged over a 4-hour period (Fig. 4G).

## DISCUSSION

In this study, we found that PS externalization is a predominant feature across different populations of small EVs in murine and human blood, including TSPN^+^ EV subsets, which have been previously described as partially or fully lacking external PS exposure (Groß et al., 2024; Lai et al., 2016; Shao et al., 2018). With the mean plasma EV size of 150 nm being the EVs analyzed here were significantly smaller than apoptotic or necrotic bodies (1-5 µm; Zhang et al., 2018). Additionally, we demonstrated that the detected PS^+^ vesicles were not primarily derived from major lipoprotein or chylomicron contaminants. Hence, exposed PS can be considered a universal EV marker. A marker that reliably identifies all EV populations is crucial for standardized EV quantification, facilitates the monitoring of EV levels for diagnostic purposes, enhances the characterization of therapeutic EVs, enables the measurement of EV turnover and biodistribution, and deepens our understanding of EV biology.

However, since PS is not exclusive to EVs, comprehensive EV characterization remains essential. While further research is needed to clarify the mechanisms of PS externalization on EV subtypes aside from apoptotic bodies and platelet-derived EVs, it is plausible that reduced flippase activity, likely from limited ATP production, contributes to PS externalization in most EVs. Additionally, localized PS exposure has been suggested to drive microvesicle formation through membrane curvature changes and increased TMEM16F activity (Wu et al., 2020; Xu et al., 2013; Benedikter et al., 2018; Kira et al., 2023; Shao et al., 2018).

The question of PS exposure on EVs has been a point of contention in the field, with studies reporting PS as characteristic of only select EV subpopulations (Heijnen et al., 1999; Jeppesen et al., 2023; Zargarian et al., 2017), while others suggest that PS exposure is more widespread (Groß et al., 2024; Matsumoto et al., 2021). Here, we demonstrate that variability in PS-binding protein affinity likely contributes to these discrepancies. Distinct affinities of Annexin V and MFG-E8 for PS have been observed in previous studies, primarily in the context of platelets (Albanyan et al., 2009; Connor et al., 2010); our study extends this comparison to liposomes, EVs, and cells. With EV flow cytometry, where low epitope density on EVs can complicate analysis, highly sensitive reagents are essential for accurate results (Welsh et al., 2020).

We observed that Annexin V exhibited low to negligible binding affinity for particles with low PS content, such as submicron-sized liposomes (1-5% PS) and blood EVs, which have been described to contain PS at low frequencies (Groß et al., 2024). However, Annexin V effectively bound to PS on necrotic cells and large liposomes (1000 nm) with high PS content (50%), indicating that its affinity may depend on both PS density and particle size or membrane curvature. Our findings are consistent with previous reports indicating that Annexin V’s anticoagulant function is reduced with PS levels below 15% (Shi et al., 2004). In contrast, MFG-E8 demonstrated higher PS affinity, especially on EVs, while the C1-Tetramer showed the highest sensitivity across EVs and cells. Given the variability in PS frequencies across EV sources, further analysis on EVs with high PS levels (e.g., from breast milk or saliva) could help clarify binding characteristics. Additionally, our findings indicate that binding affinities for MFG-E8 and C1-Tetramer are influenced by both particle size and PS density, with dissociation constants ranging from 4.5 to 96 nM, consistent with previously published data for full-length MFG-E8 (Borisenko et al., 2004; Shao et al., 2008; Suwatthee et al., 2023). A full understanding of factors influencing MFG-E8 and C1-Tetramer binding affinities remains beyond this study, but careful titration of PS-binding proteins is recommended for EV analysis.

Leveraging the prevalence of PS on plasma EVs, we used MFG-E8-based reagents to study *in vivo* EV turnover. While prior studies have investigated EV clearance by reintroducing exogenously labeled EVs (Kang et al., 2021) or using genetic markers to trace EV subpopulations (Boudna et al., 2024; Verweij et al., 2021), our study offers a novel approach to track endogenous EV turnover. Exogenous EVs are cleared rapidly from circulation (95% within 5 min; Matsumoto et al., 2017; Morishita et al., 2015; Takahashi et al., 2013), whereas our data suggest a slower clearance rate for endogenous EVs (t_1/2_ = 29 min). This discrepancy might stem from differences in PS frequency or from PS-independent clearance mechanisms. To our knowledge, this study is the first to employ broad *in vivo* labeling, enabling insights into the turnover of circulating and cell-bound endogenous EVs in physiological and pathological conditions. Future research could elucidate whether reductions in serum EVs observed during acute infections, such as LCMV, are linked to altered turnover rates (Rausch et al., 2023).

Although we cannot entirely exclude the possibility that C1-Tetramer labeling affects EV clearance, we observed no reduction in PS exposure on labeled EVs, evidenced by consistent *in vivo* and *in vitro* PS co-staining. This finding is relevant given PS’s known role in EV uptake via multiple receptors (Brodeur et al., 2023; Buzas et al., 2018; Mulcahy et al., 2014), including Tim4, RAGE, Bai-1, and stabilin-2, all of which are expressed on macrophages essential for PS^+^ EV clearance (Matsumoto et al., 2017; Matsumoto et al., 2021). Consistent with these observations, we found that PS^+^ EVs predominantly interact with CD11b^+^ monocytes/macrophages and CD19^+^ B cells in the spleen (Kranich et al., 2020). Furthermore, our results showed that after clearance from circulation, endogenously labeled PS^+^ EVs persist on spleen cell surfaces but are also internalized by macrophages. This suggests that splenic EV retention outlasts their circulation time, likely due to interactions with PS receptors on target cells. Studying EV kinetics in other tissues, particularly the liver - an organ known for macrophage-mediated EV clearance - could provide further insights (Matsumoto et al., 2021).

## ACKNOWLEDGEMENTS

We acknowledge the Core Facility Flow Cytometry at the Biomedical Center, Ludwig Maximilian University of Munich, for providing the ImageStreamX MKII imaging flow cytometer, cell sorters, and services. T.B., A.K. and J.K are supported by the Deutsche Forschungsgemeinschaft (German Research Foundation)—Project-ID 210592381–SFB 1054 (TP A06, B03, Z02).

