## Supplemental data for "Assessing Extracellular Vesicle Turnover In vivo Using Highly Sensitive Phosphatidylserine-Binding Reagents"

### Supplementary Figures

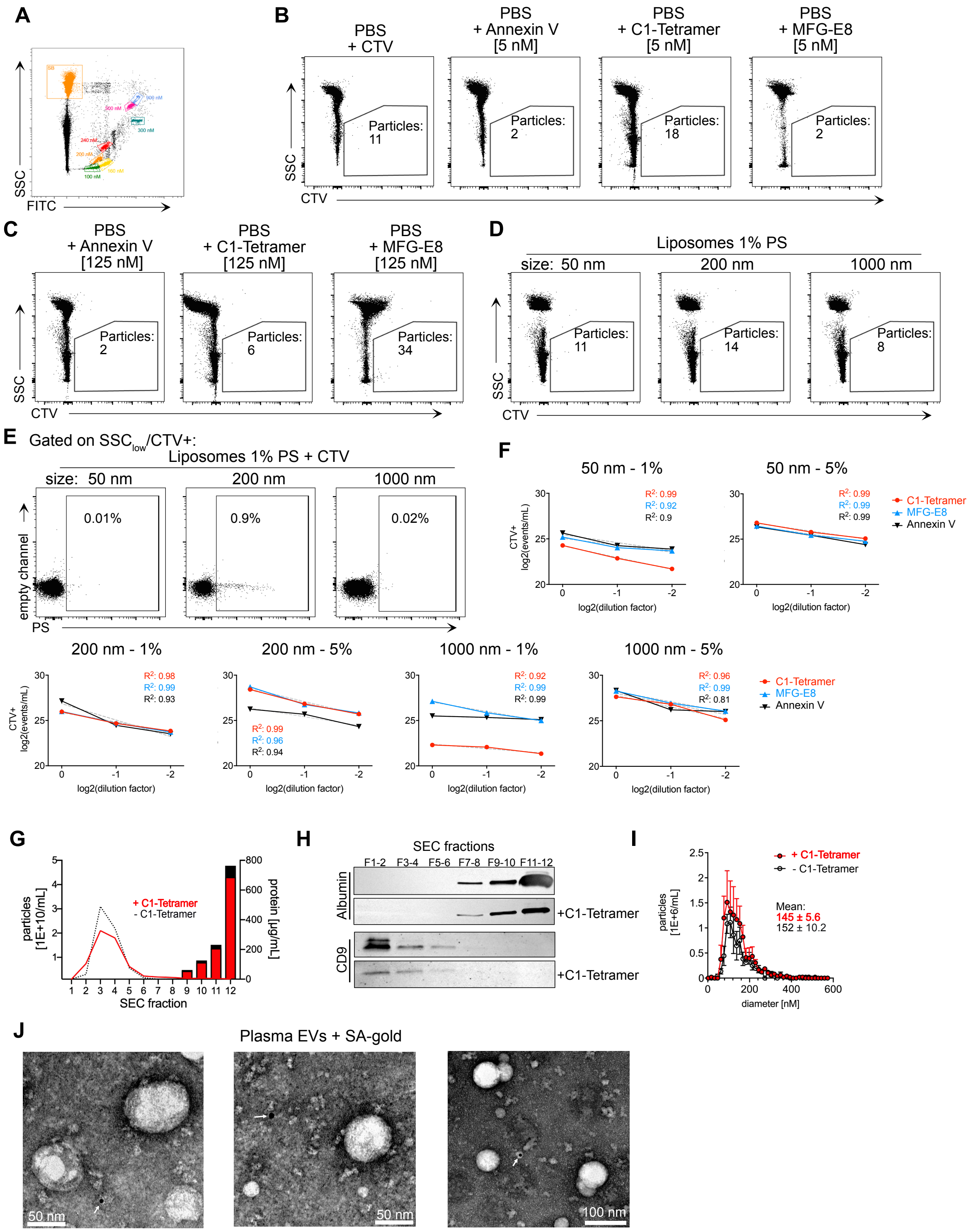


**Suppl. Fig. 1: (A)** Megamix-Plus (BioCytex) SSC and FSC beads with distinct submicron sizes analyzed by imaging flow cytometry (IFC), all bead populations, ranging from 100 nm to 900 nm could be resolved based on scatter and fluorescent emission in Ch02 on the ImageStream$^{TM}$. SB, speed beads. **(B-E)**, Imagestream$^{TM}$ analysis of liposomes (Encapsula), shown are exemplary graphs for aggregate controls at 5 nM and 125 nM of all staining reagents in **(B)** and **(C)**, respectively, unstained liposomes are depicted in **(D)** and PS FMO liposomes stained only with CTV are shown in **(E)**. **(F)**, Exemplary twofold serial dilution for all liposome populations stained with CTV and either Annexin V (FITC), MFG-E8-eGFP or the MFG-E8 C1-Tetramer (AF488) at 5 nM, measured by IFC. Linear regression analysis (grey dashed lines) was performed to check for swarm detection, goodness of fit is indicated by R-square values. **(G)** Elution profile of murine plasma EVs unlabelled or labelled with the C1-Tetrarmer prior to size exlcusion chromatography (SEC), briefly, EVs were isolated via SEC and the collected fractions were analysed by nanoparticle tracking analysis (NTA) for particle concentration/mL (lines) and by bicinchoninic acid assay (BCA) to determine the protein content in each fraction (bars). **(H)** Western blot of SEC fractions for negative control serum albumin and the EV marker CD9 as a positive control. **(I)** Size distribution analysis of pooled fractions (F1-5) for unlabelled vs. labelled plasma EVs. **(J)** Transmission electron microscopy images of murine plasma EVs incubated with SA-gold as a negative control. White arrows indicate gold particles.

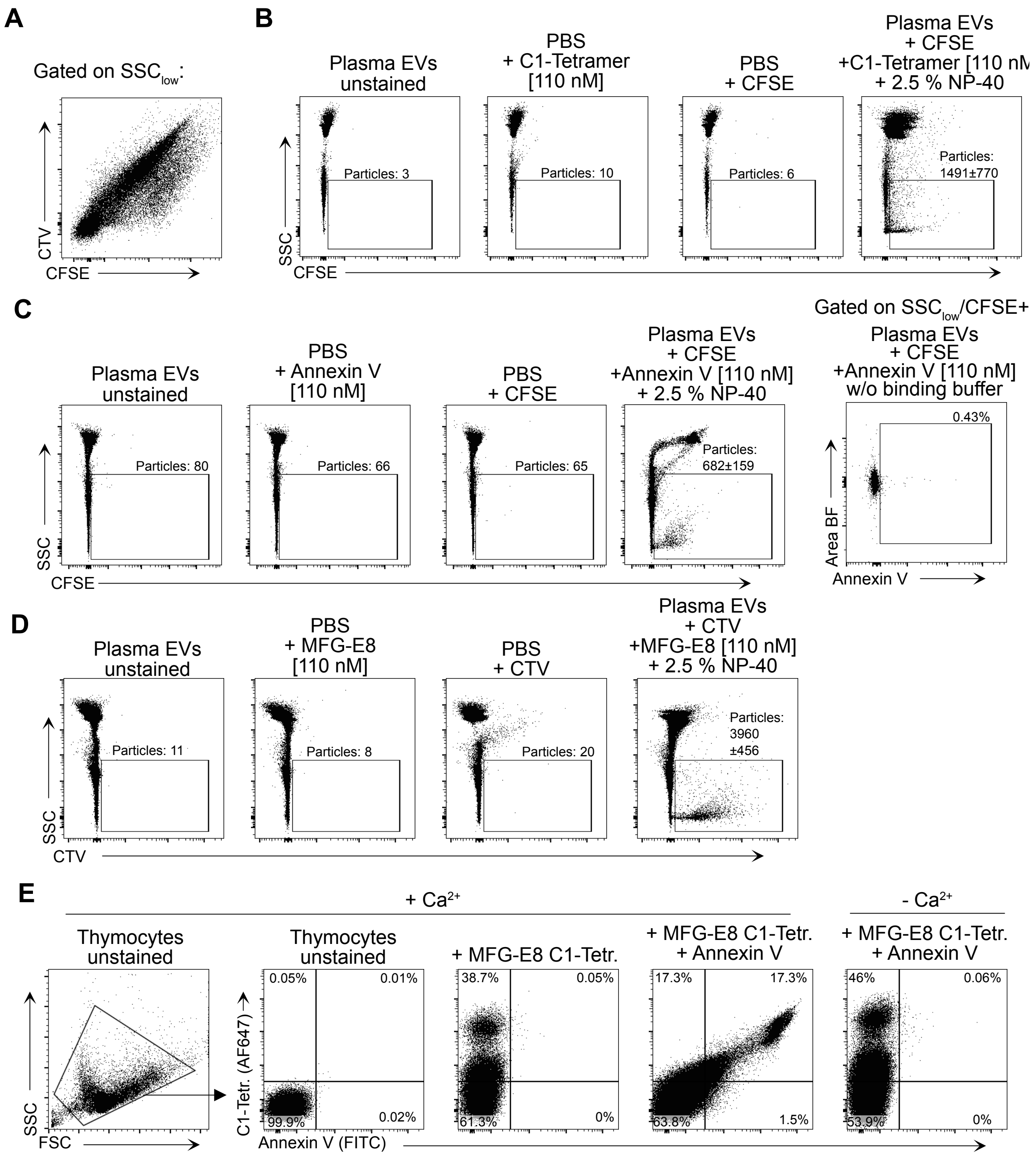


**Suppl. Fig. 2: (A)** Co-staining of membrane dyes, CTV and CFSE, used for IFC analysis of plasma EVs. Exemplary graphs shown for controls of plasma EV IFC analysis including unstained EVs, detergent controls and aggregates controls for murine plasma EVs labelled with 110 nM of **(B)** C1-Tetramer (AF647) **(C)** Annexin V (AF647) or **(D)** MFG-E8-eGFP (main Fig. 1H). **(E)**, Staurosporine-treated (1 µg/ml, 3 h) apoptotic murine (C56BL/6 mice) thymocytes were stained either with Annexin V or MFG-E8 C1-Tetramer or both reagents together in $Ca^{2+}$-containing or $Ca^{2+}$-free buffer as indicated and analyzed by flow cytometry.

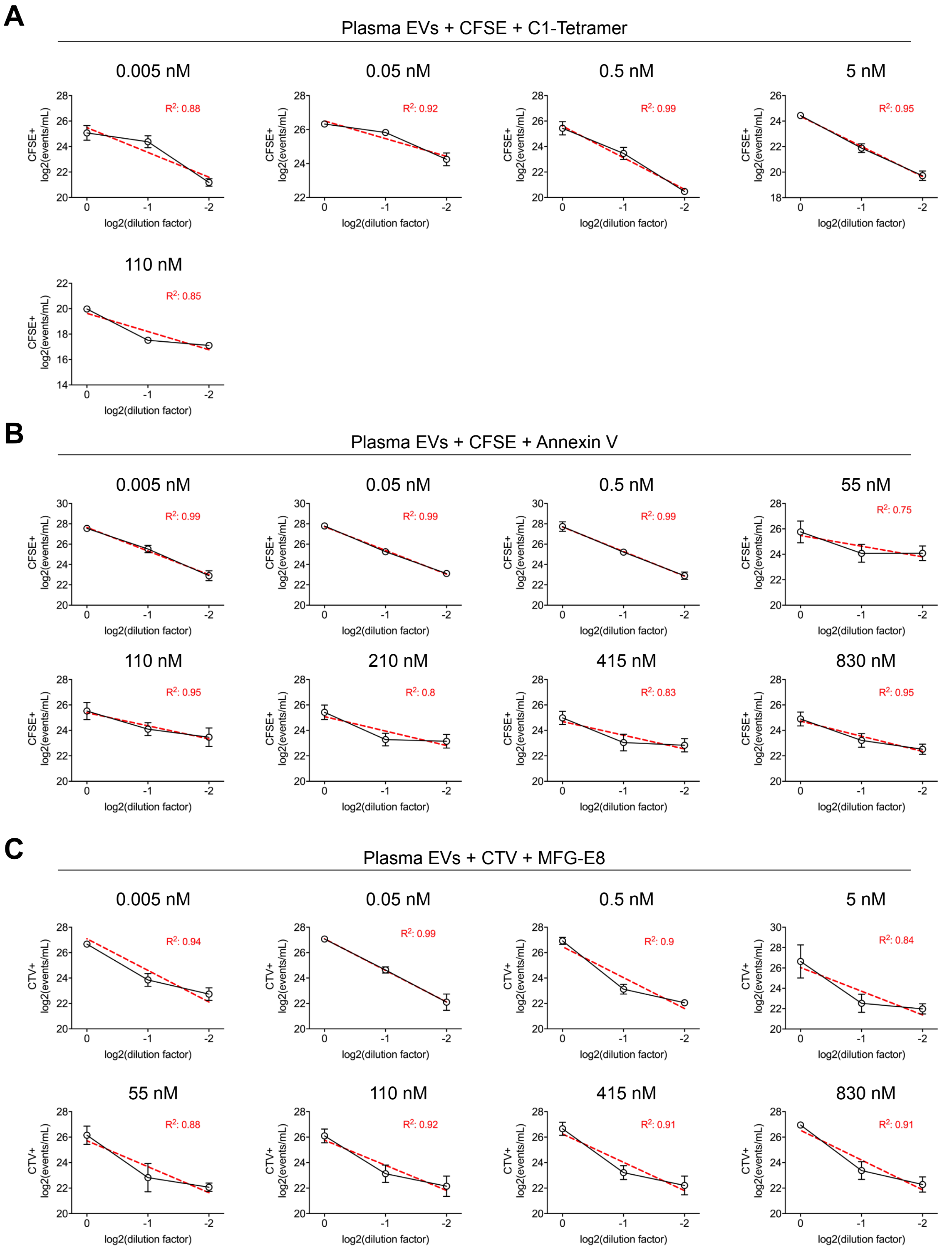


**Suppl. Fig. 3: (A-C)** Twofold serial dilution of plasma EVs (main Figure 1 G,H) stained with CFSE or CTV and different concentrations of **(A)** C1-Tetramer (AF647), **(B)** Annexin V (AF647) **(C)** MFG-E8-eGFP (n=3), measured by IFC. Linear regression analysis (red dashed line) was performed for each dilution series to check for swarm detection, goodness of fit is indicated by R-square values.

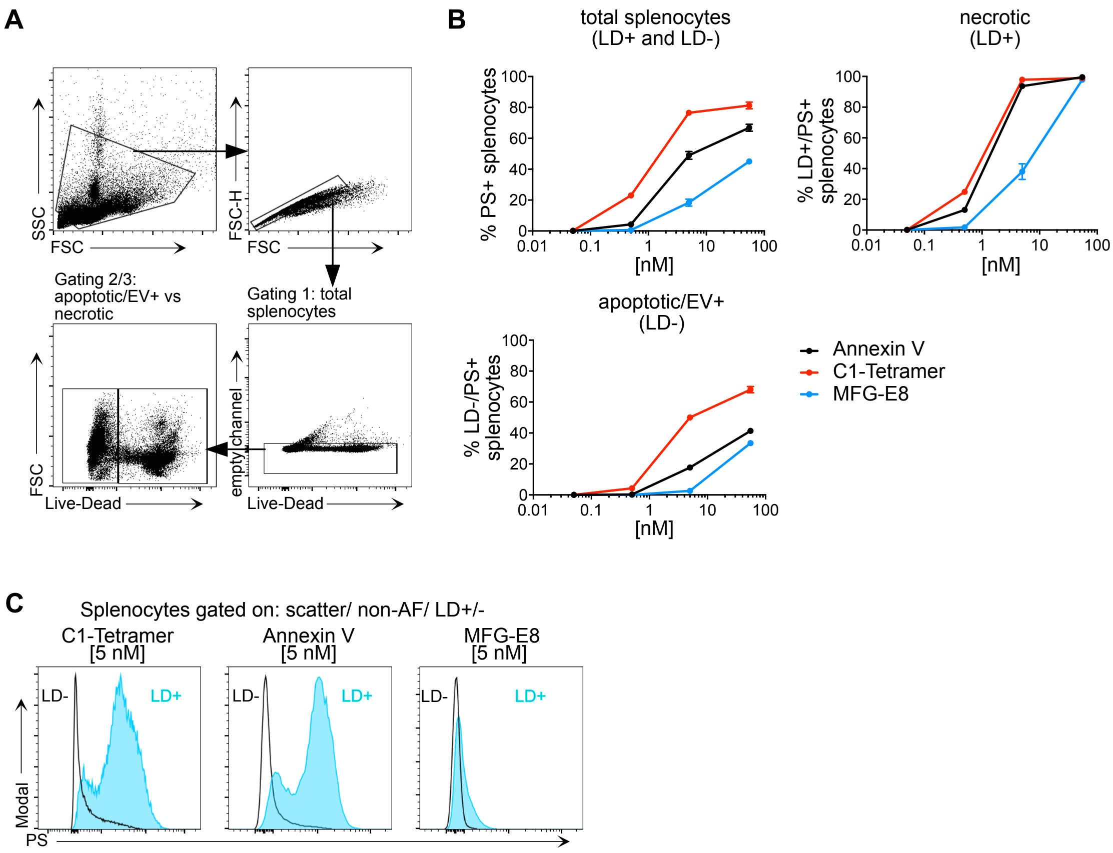


**Suppl. Fig. 4: (A)** Gating strategy for splenocytes analysed by flow cytometry. Briefly, cells were heat-shocked for 3 min at 60°C to induce cell death and increase PS exposure. **(B)** PS-labelling of total, necrotic (LD+) or apoptotic/EV+ (LD-) splenocytes using distinct concentrations of Annexin V-FITC (black), C1-Tetramer-AF647 (red) or MFG-E8-eGFP (blue) (n=3). **(C)** Histograms for PS fluorescent intensities of Live-Dead (LD) + vs - splenocytes stained with different PS-binding reagents.

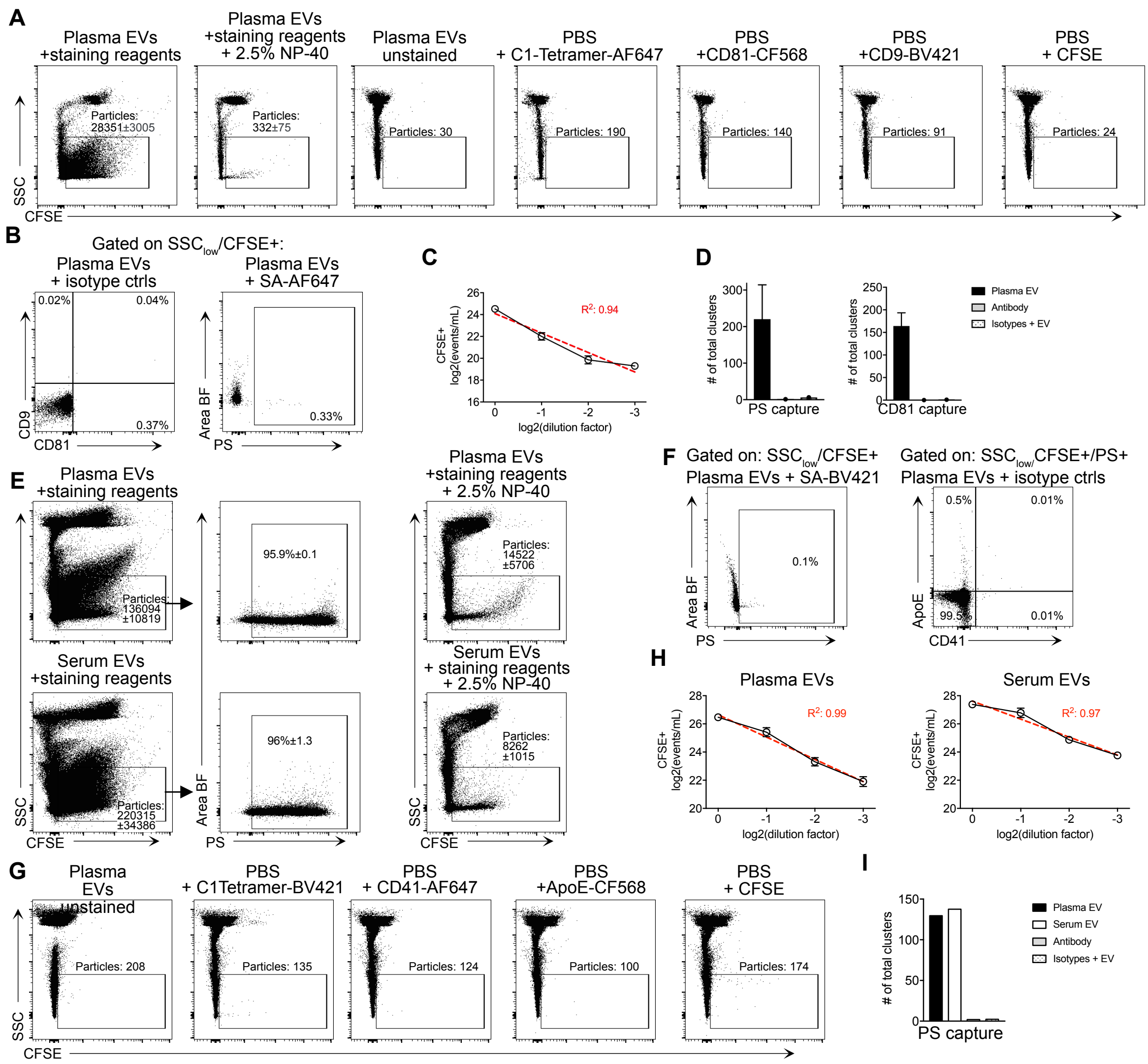


**Suppl. Fig. 5: (A)** Gating on SSClow/CFSE+ events for plasma EVs (main Fig. 2A) and detergent controls (n=3) as well as unstained EVs and aggregate controls for all staining reagents. **(B)** Matched isotype controls for CD9 and CD81 antibodies and SA-AF647 staining as a negative control for the C1-Tetramer-AF647 **(C)** Twofold serial dilution of plasma EVs stained with CFSE, C1-Tetramer, CD9 and CD81, measured by IFC. Linear regression analysis (red dashed line) was performed to check for swarm detection, goodness of fit is indicated by R-square values. **(D)** Antibody only and matched isotype controls for dSTORM analysis of murine EVs using different capturing reagents (main Fig. 2C), shown is the number of total clusters as determined by cluster analysis. **(E)** Gating on SSClow/CFSE+ and PS+ murine serum or plasma EVs and detergent controls (n=3) for main Fig. 2D. **F** Matched isotype controls for CD41 and ApoE antibodies and SA-BV421 staining as a negative control for the C1-Tetramer-BV421. **(G)** Gating on SSClow/CFSE+ events for unstained EVs and aggregate controls for all staining reagents used in main Figure 2D. **(H)** Twofold serial dilution of plasma EVs stained with CFSE, C1-Tetramer, CD41 and ApoE, measured by IFC. Linear regression analysis (red dashed line) was performed to check for swarm detection, goodness of fit is indicated by R-square values. **(I)** Antibody only and matched isotype controls for dSTORM analysis of murine PS+ EVs (main Fig. 2E), shown is the number of total clusters as determined by cluster analysis.

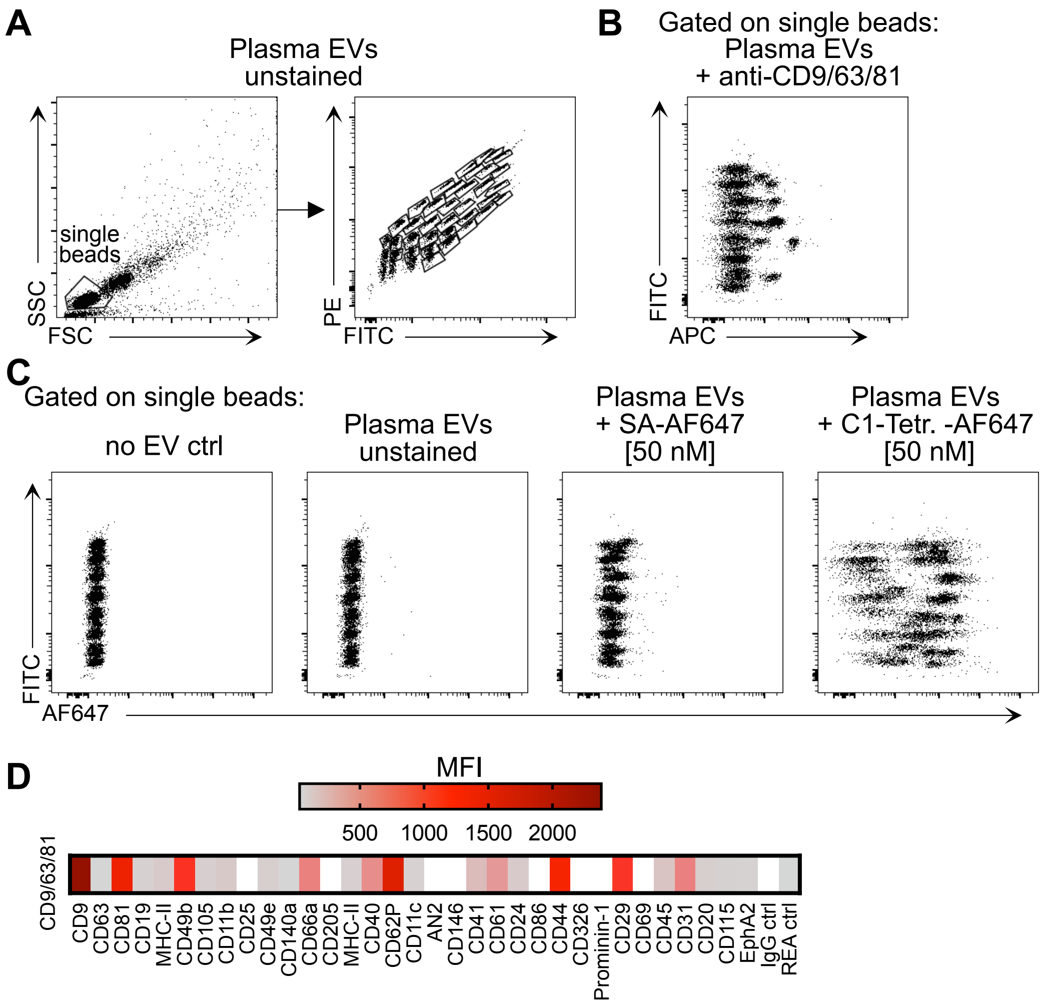


**Suppl. Fig. 6: (A)** Exemplary gating strategy for bead-assisted flow cytometry as shown in Figure 2F. Single beads are first gated by SSC/FSC-A and individual beads containing antibodies directed against typical EV surface markers then separated by their endogenous PE/FITC-signal, enabling discrimination of 35 bead populations. **(B)** Detection antibody mix (Miltenyi) was used as an internal assay control.**(C)** AF647 fluorescent intensity of background controls ('no EV ctrl', 'plasma EVs unstained' and 'plasma EVs + SA-AF647') versus plasma EVs detected by C1-Tetramer-AF647 labelling. **(D)** Surface exposure of CD9/CD63/CD81 (row) analyzed by bead-assisted flow cytometry, using a detection antibody mix (Miltenyi). Distinct EV subpopulations are shown which were captured by bead-coupled antibodies targeting the indicated proteins (columns). Color coding indicates MFI for APC of the respective bead population, MFI values were background corrected by a 'no EV' blank control.

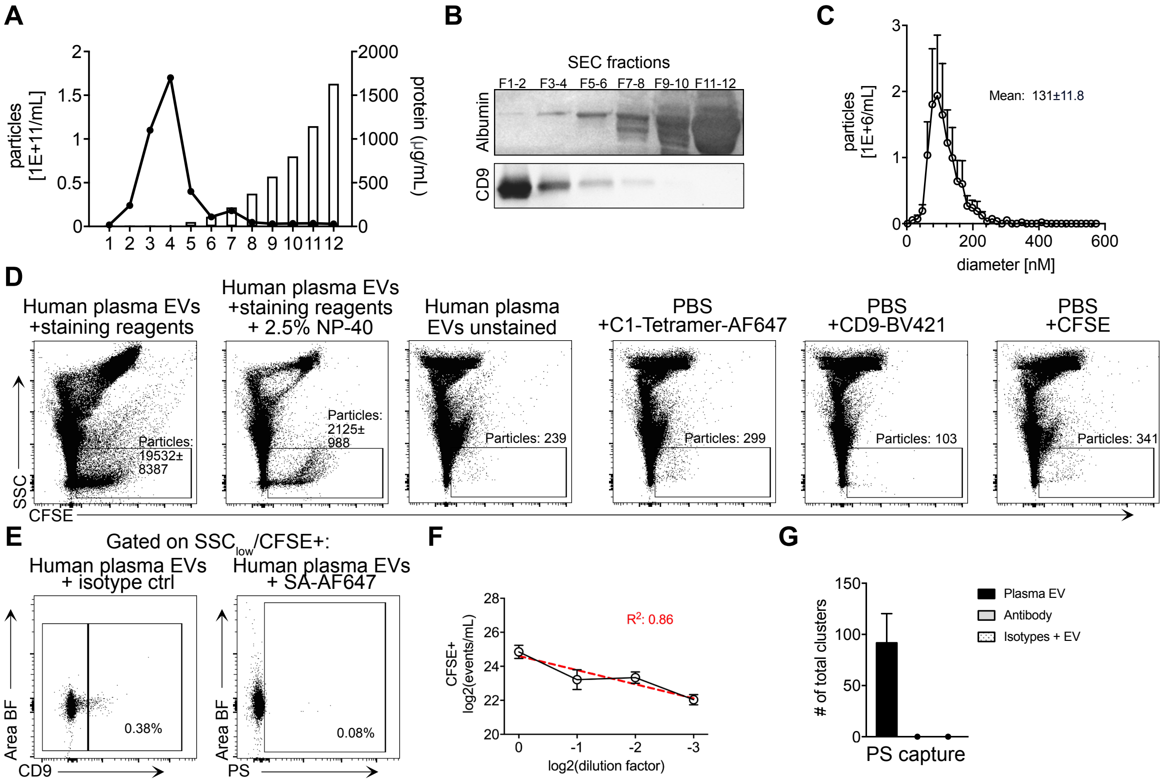


**Suppl. Fig. 7: (A)** Elution profile of human plasma EVs, briefly, EVs were isolated via SEC and the collected fractions were analysed by nanoparticle tracking analysis (NTA) for particle concentration/mL (lines) and by bicinchoninic acid assay (BCA) to determine protein content in each fraction (bars). **(B)** Western blot of SEC fractions for negative control serum albumin and the EV marker CD9 as a positive control. **(C)** Size distribution analysis of pooled fractions (F1-5). **(D)** Gating on $SSC_{low}$/CFSE+ events for human plasma EVs (main Fig. 3A) and detergent controls (n=5) as well as unstained EVs and aggregate controls for all staining reagents. **(E)** Matched isotype control for CD9 and SA-AF647 as a negative control for the C1-Tetramer **(F)** Twofold serial dilution of human plasma EVs stained with CFSE, C1-Tetramer and CD9, measured by IFC. Linear regression analysis (red dashed line) was performed to check for swarm detection, goodness of fit is indicated by R-square values.**(G)** Antibody only and matched isotype control for dSTORM analysis of human EVs (main Fig. 3B), shown is the number of total clusters as determined by cluster analysis.

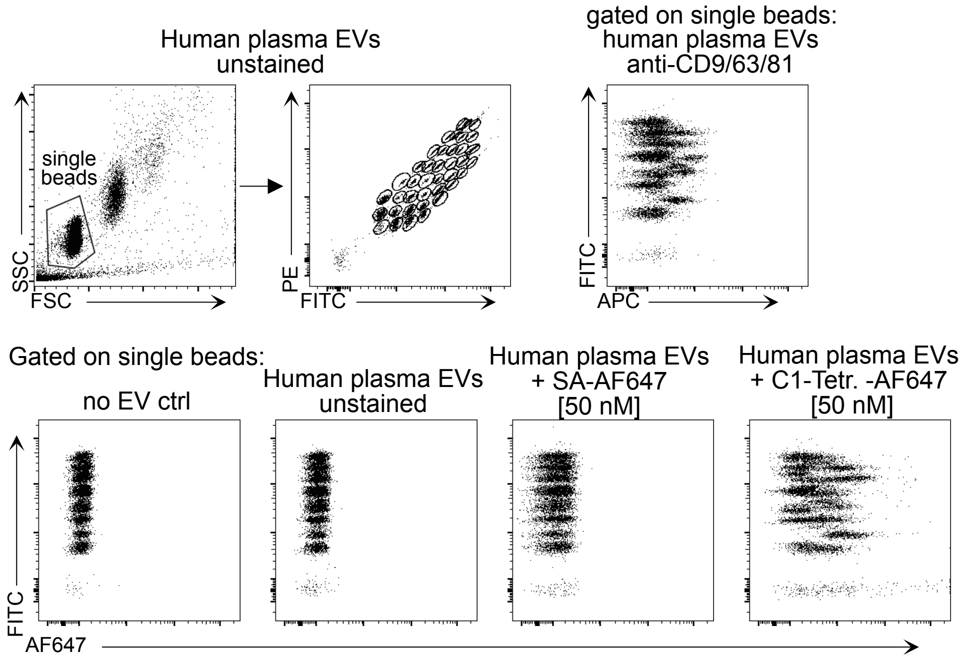


**Suppl. Fig. 8: (A)** Exemplary gating strategy for bead-assisted flow cytometry as shown in main Fig. 3C. Single beads are first gated by SSC/FSC-A and individual beads containing antibodies directed against typical EV surface markers then separated by their endogenous PE/FITC-signal, enabling discrimination of 39 bead populations. **(B)** Detection antibody mix (Miltenyi) was used as an internal assay control. **(C)** AF647 fluorescent intensity of background controls ('no EV ctrl', 'plasma EVs unstained' and 'plasma EVs + SA-AF647') versus plasma EVs detected by C1-Tetramer-AF647 labelling.

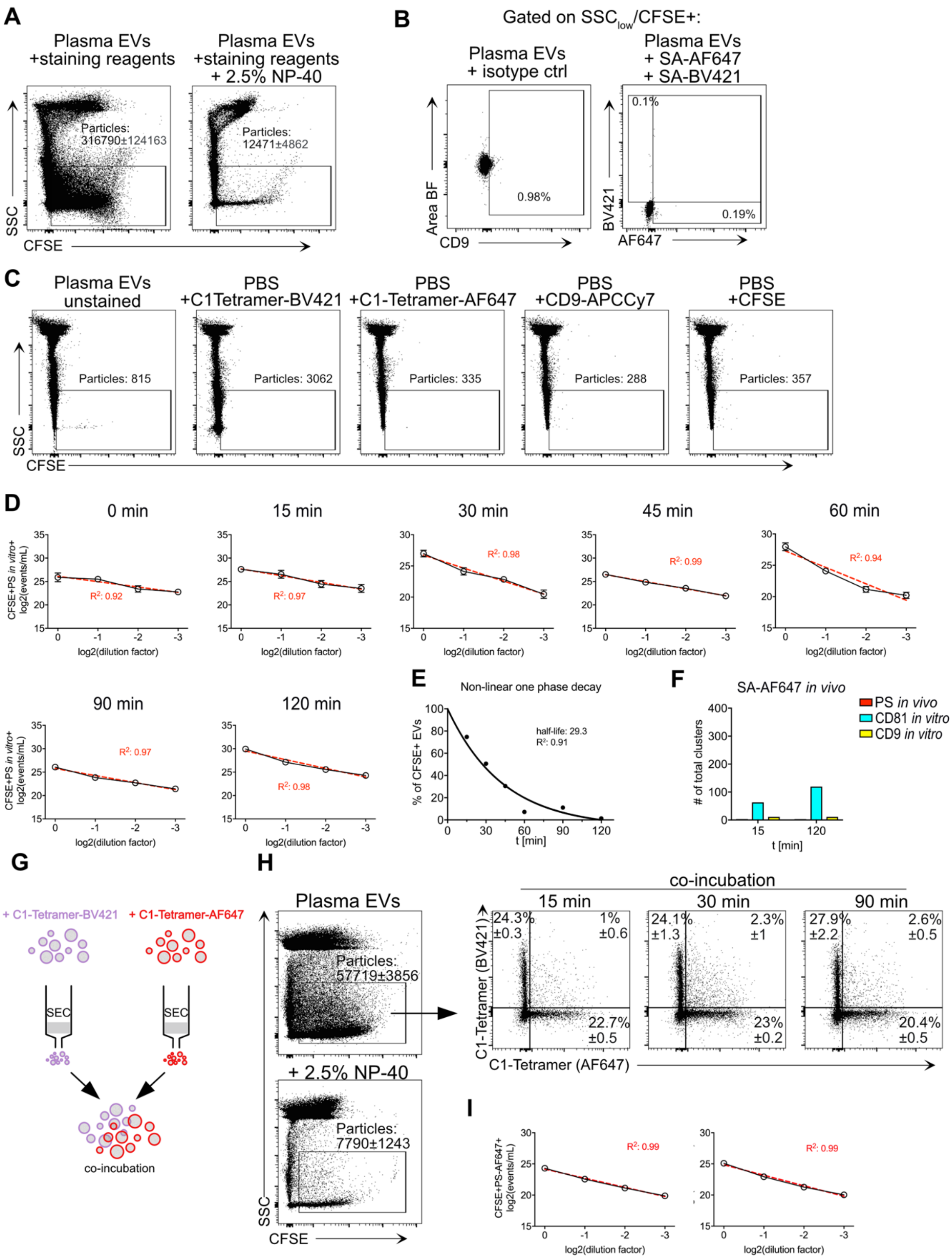


**Suppl. Fig. 9: (A)** Gating on $SSC_{low}$/CFSE+ events for *in vivo* labelled plasma EVs (t=15 min, n=5)(main Fig. 4B) **(B)** Matched isotype control for CD9 and SA-BV421 staining as a negative control for the C1-Tetramer *in vitro* staining. **(C)** Exemplary graphs after 15 min of C1-Tetramer i.v. injection, showing detergent controls (n=5) as well as unstained EVs and aggregate controls for all staining reagents. **(D)** Twofold serial dilution of *in vivo* PS-labelled plasma EVs stained with CFSE, C1-Tetramer *in vitro/in vivo* and CD9 measured by IFC. Linear regression analysis (red dashed line) was performed to check for swarm detection, goodness of fit is indicated by R-square values. **(E)** Half-live of in vivo labelled plasma EVs (main Fig. 4C) was determined by fitting a non-linear one phase decay curve. **(F)** EV clusters at two timepoints (15 and 120 min) after i.v. injection of SA-AF647. Bar graphs show number of *in vivo* stained SA+ (red), *in vitro* stained CD81+ (cyan) and CD9+ (yellow) EV clusters. **(G,H)** To assess stability of the C1-Tetramer EV labelling, separate plasma EV samples were stained with C1-Tetramer in BV421 or AF647. Afterwards free dye was removed by SEC and the distinctly labelled EVs were pooled and incubated at 36°C for 15, 30 or 90 min (n=3). To check for dye transfer, EVs were analyzed by IFC. **(I)** Twofold serial dilutions and linear regression analysis (red dashed line) was performed for the C1-Tetramer labelled EVs to control for swarm detection. Goodness of the fit is indicated as R-square values.

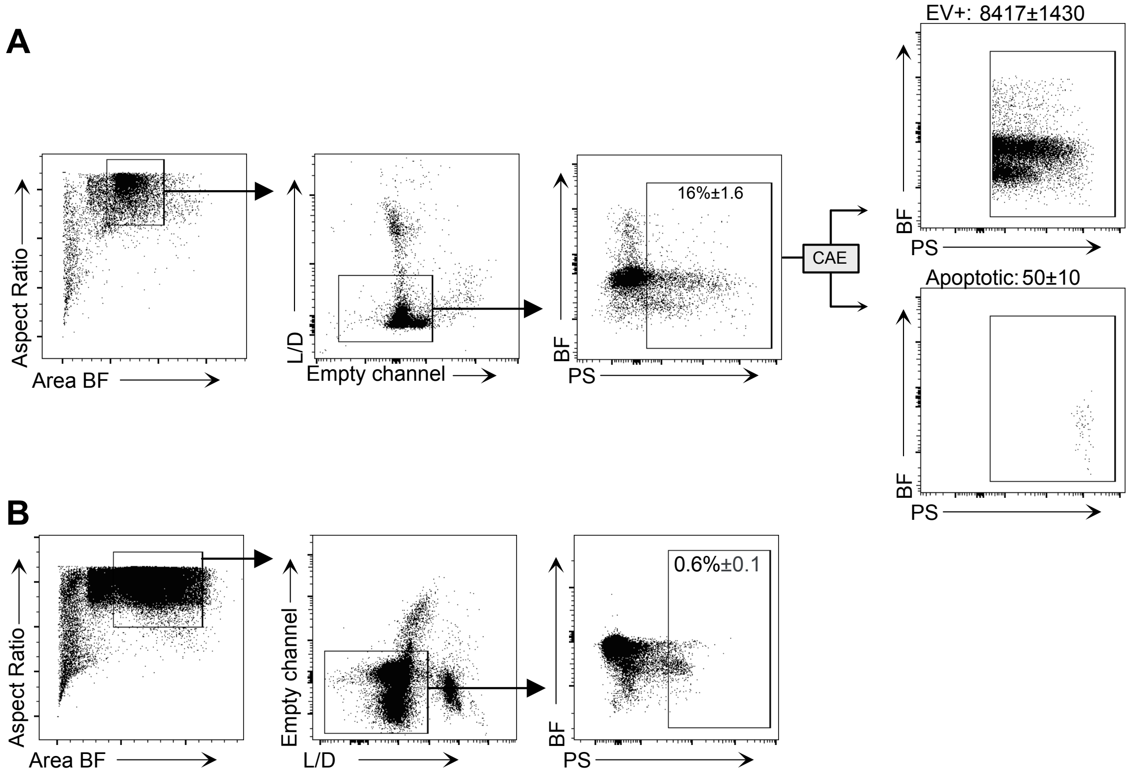


**Suppl. Fig. 10: (A)** To discriminate dying and EV+ cells, murine splenocytes were analysed using IDEAS, a machine-learning based convolutional autoencoder (CAE) and FlowJo, as described previously (Kranich et al., 2020; Rausch et al., 2023; Rausch et al., 2021).**(B)** Control mice were injected with 50 µg SA-AF647 (n=3) and splenocytes were analysed by IFC after 15 min, to check for unspecific labelling.
