## Supplemental tables for "Assessing Extracellular Vesicle Turnover In vivo Using Highly Sensitive Phosphatidylserine-Binding Reagents"

**Supplementary Table S1.** Framework representing the Minimal Information about a Flow Cytometry (FC) experiment for EV-FC-specific reporting (MIFlowCyt-EV template), as recommended by the Minimum Information for Studies of EVs (MISEV).

| Framework Criteria | What to report | Please complete each criterion |
|---|---|---|
| 1.1 Preanalytical variables conforming to MISEV guidelines. | Preanalytical variables relating to EV sample including source, collection, isolation, storage, and any others relevant and available in the performed study. | All mice were housed under specific pathogen-free conditions at the Core Facility Animal Models of the Biomedical Center (Ludwig-Maximilians-University, Munich). Blood from 6-8 weeks old female C57BL/6JNRj mice (Janvier) was collected by heart puncture. For serum, blood samples were allowed to coagulate for 30 min at RT whereas heparin was added to plasma samples. Subsequently, blood was centrifuged at 1,000 g for 10 min at 4°C and serum or plasma was diluted 1:1 with 1X cOmplete™ protease inhibitor cocktail (Roche). Alternatively, aliquots of plasma or serum were snap-frozen and stored at -20°C at this step. Samples were centrifuged sequentially at 2,000 g at 4°C and 10,000 g at RT for 10 min. Afterwards, 800-900 µL of sample were applied onto SEC columns (qEV1, Gen2 Izon) and EV-containing fractions were collected and concentrated to 300 µL. EV samples were not stored prior to the measurement but used freshly. Human plasma was collected from 5 healthy individuals (3 males, 2 females – age 21-60) into S-Monovette® Natrium Heparin tubes and processed as described above for murine samples. |
| 1.2 Experimental design according to MIFlowCyt guidelines. | EV-FC manuscripts should provide a brief description of the experimental aim, keywords, and variables for the performed FC experiment(s) using MIFlowCyt checklist criteria: 1.1, 1.2, and 1.3, respectively. Template found at <u>www.evflowcytometry.org.</u> | **1.1** Aim: Determine the frequency of PS among blood EVs and analyze circulating and cell-bound PS+ EVs to investigate their clearance *in vivo* **1.2** Phosphatidylserine; Extracellular Vesicles; Imaging flow cytometry; superresolution microscopy; clearance **1.3** Experimental variables: a) murine plasma/serum from 6-8 weeks old female C57BL/6J mice were stained with CFDA-SE, CellTrace Violet, CD9, CD81, CD41, ApoE, C1-Tetramer (MFG-E8 derivative), Annexin V and/or MFG-E8-eGFP and measured by Imaging Flow Cytometry (and superresolution microscopy). b) Human plasma EVs from 5 healthy individuals were stained with CFDA-SE, CD9 and C1-Tetramer (MFG-E8 derivative) and measured by Imaging Flow Cytometry (and superresolution microscopy). |
| 2.1 Sample staining details | State any steps relating to the staining of samples. Along with the method used for staining, provide relevant reagent descriptions as listed in MIFlowCyt guidelines (Section 2.4 Fluorescence Reagent(s) Descriptions). | **Anti-mouse mAB:**<br>• Anti-mouse CD9 eFluor450 (clone: KMC8. isotype: rat IgG2a) (eBiosciences) (final concentration for staining in 60 µL: 120 ng/test)<br>• Anti-mouse CD9 APC/Fire™ 750 (clone: MZ3. isotype: rat IgG2a) (Biolegend) (final concentration for staining in 60 µL: 120 ng/test)<br>• Purified anti-mouse CD81 (clone: Eat-2. Isotype: Armenian Hamster IgG1k) fluorescently labelled with Mix-n-Stain CF568 Antibody labelling kit (Sigma-Aldrich) (final concentration for staining in 60 µL: 300 ng/test)<br>• Purified anti-mouse CD41 (clone: MWReg30. Isotype: rat IgG1k) (Biolegend) fluorescently labelled with Mix-n-Stain CF647 antibody labelling kit (Sigma-Aldrich) (final concentration for staining in 60 µL: 300 ng/test) |

| | | |
|---|---|---|
| | | • Purified anti-mouse ApoE (clone: 3B3C32. Isotype: rat IgG2a,k) (Biolegend) labelled with Mix-n-Stain CF568 antibody labelling kit (Sigma-Aldrich) (final concentration for staining in 60 µL: 300 ng/test)<br><br>**Anti-human mAB:**<br>• Anti-human CD9 BV421 (clone: M-L13. Isotype: mouse IgG1k) (BD biosciences) (final concentration for staining in 60 µL: 120 ng/test)<br><br>**Membrane dyes:**<br>• Carboxyfluorescein succinimidyl ester (CFSE, eBiosciences) (final concentration in 60 µL: 15 µM)<br>• CellTrace Violet (Thermo Fisher Scienitific) (final concentration in 60 µL: 15 µM)<br><br>**PS-binding reagents:**<br>• C1-Tetramer (MFG-E8 biotinylated monomeric C1-domain tetramerized)(unless stated otherwise final concentration was 5 nM in 60 µL)<br>• Annexin V FITC or AF647 (Biolegend) (final concentration varied from 0.005-830 nM in 60 µL)<br>• Recombinant full-length MFG-E8-eGFP (Kranich *et al.*, 2020)(final concentration varied from 0.005-830 nM in 60 µL)<br><br>**PS-binding reagents: Annexin V and C1-Tetramer. Staining preparation:** All staining reagents were centrifuged at 18,000g for 30 min at 4°C. 1 µL of CFSE or CTV were added to 40 µL of sample material to reach a final concentration of 10 µM. After addition of the membrane dyes, samples were incubated for 1 h at RT protected from light. Antibodies or staining reagents were added (or from predilutions) to the samples to reach the final concentrations as stated above in a total volume of 60 µL (to reach 60 µL among all samples spare volumes of filtered PBS were added if necessary). For Annexin V staining, 6 µL of 10X Annexin binding buffer were added to the samples while the total volume of 60 µL remained the same among all samples. Subsequently, staining was done for another hour at RT. Afterwards, samples were split in half and 2.5% (v/v) NP-40 was added to detergent controls whereas the same volume of PBS was added to the samples. Prior to IFC measurements, samples were diluted to 120 µL with filtered PBS or 120 µL filtered PBS with 1X Annexin V binding buffer. Aggregate controls contained the same final concentration of antibodies, staining reagents or membrane dyes diluted in filtered PBS. |
| 2.2 Sample washing details | State any steps relating to the washing of samples. | No washing steps were included in our protocols. Background fluorescence was accounted for by fluorescence-minus-one controls, antibody buffer, buffer only and unstained EV samples. |

| | | |
|---|---|---|
| 2.3 Sample dilution details | All methods and steps relating to sample dilution. | Samples were diluted 1:3 in filtered PBS before measurements. For serial dilutions, samples were diluted three to four times (two-fold each step). |
| 3.1 Buffer alone controls. | State whether a buffer-only control was analyzed at the same settings and during the same experiment as the samples of interest. If utilized it is recommended that all samples be recorded for a consistent set period of time e.g. 5 minutes, rather than stopping analysis at a set recorded event count e.g. 100,000 events. This allows comparisons of total particle counts between controls and samples. | Buffer only controls were analyzed at the same settings and at least once for each experimental condition at the same time as the experiment itself. All samples were measured for at least 1 minute. |
| 3.2 Buffer with reagent controls. | State whether a buffer with reagent control was analyzed at the same settings, same concentrations, and during the same experiment as the samples of interest. If used state what the results were. | Buffer with reagent controls ("aggregate controls") were acquired for all staining reagents used in the respective experiment. Reagents were used at the same concentrations as used for staining and stated in section 2.1. All buffers with reagent controls were analyzed at the same settings and at least once for each experimental condition at the same time as the experiment itself. All samples were measured for at least 1 minute. Exemplary results of particles detected as $SSC_{low}$/CFSE or CTV+ events can be found in the supplementary figures. |
| 3.3 Unstained controls. | State whether unstained control samples were analyzed at the same settings and during the same experiment as stained samples. If used, state what the results were, preferably in standard units. | Unstained control samples were analyzed at the same settings and included in all experiments. Exemplary results of particles detected as $SSC_{low}$/CFSE or $SSC_{low}$/CTV+ events can be found in the supplementary figures or are also available upon request. |
| 3.4 Isotype controls. | The use of isotype controls is applicable to immunofluorescence labelling only. State whether isotype controls were analyzed at the same settings and during the same experiment as stained | **Anti-mouse mAB matched isotype controls:**<br>• For anti-mouse CD9 eFluor450 (eBiosciences) the following isotype control antibody was used: Rat IgG2a,l isotype control antibody BV421 (clone: RTK2759, Biolegend) (final concentration for staining in 60 µL: 120 ng/test)<br>• For anti-mouse CD9 APC/Fire$^{TM}$ 750 (clone: MZ3. isotype: rat IgG2a) the following isotype control antibody was used: |

| | | |
|---|---|---|
| | samples. If utilized, state which antibody they are matched to, the concentration used, and what the results were (Section 4.2, 4.3, 4.4). Due to conjugation differences between manufacturers if should be stated if the isotype controls are from the same manufacturer as the matched antibodies. | Rat IgG2a,l isotype control antibody APCCy7 (clone: RTK2758, Biolegend) (final concentration for staining in 60 µL: 120 ng/test)<br>• For purified anti-mouse CD81 (clone: Eat-2. Isotype: Armenian Hamster IgG1k) fluorescently labelled with Mix-n-Stain CF568 Antibody labelling kit (Sigma-Aldrich) the following isotype control antibody was used:<br>Purified armenian hamster IgG isotype ctrl antibody (clone: HTK888, Biolegend) fluorescently labelled with Mix-n-Stain CF568 Antibody labelling kit (Sigma-Aldrich) (final concentration for staining in 60 µL: 300 ng/test)<br>• For purified anti-mouse CD41 (clone: MWReg30. Isotype: rat IgG1k) (Biolegend) fluorescently labelled with Mix-n-Stain CF647 antibody labelling kit (Sigma-Aldrich) the following isotype control antibody was used:<br>Purified rat IgG1k isotype ctrl antibody (clone: RTK2071, Biolegend) fluorescently labelled with Mix-n-Stain CF647 antibody labelling kit (Sigma-Aldrich) (final concentration for staining in 60 µL: 300 ng/test)<br>• For purified anti-mouse ApoE (clone: 3B3C32. Isotype: rat IgG2a,k) (Biolegend) labelled with Mix-n-Stain CF568 antibody labelling kit (Sigma-Aldrich) the following isotype control antibody was used:<br>Rat IgG2a,k isotype control antibody PE (clone:RTK2758) (final concentration for staining in 60 µL: 300 ng/test)<br><br>**Anti-human mAB matched isotype controls:**<br>• For Anti-human CD9 BV421 (clone: M-L13. Isotype: mouse IgG1k) (BD biosciences) the following isotype control antibody was used: mouse IgG1,k isotype control antibody BV421 (clone: MOPC-21, Biolegend) (final concentration for staining in 60 µL: 120 ng/test) |
| 3.5 Single-stained controls. | State whether single-stained controls were included. If used state whether the single-stained controls were recorded using the same settings, dilutions, and during the same experiment as stained samples and state what the results were, preferably in standard units (Section 4.2, 4.3, 4.4). | For compensation, single stained controls were prepared for each fluorophore with BD CompBeads (Anti-Mouse Ig or Anti-Hamster) and BD CompBeads Negative controls. The compensation beads were acquired at the same laser settings and antibodies were diluted as stated above for the staining. Compensation matrices were calculated using the IDEAS software. |

| | | |
|---|---|---|
| 3.6 Procedural controls. | State whether procedural controls were included. If used, state the procedure and if the procedural controls were acquired at the same settings and during the same experiment as stained samples. | Plasma EVs were used for fluorescence-minus-one controls and/or isotype controls were measured at the same settings and during each experiment. For C1-Tetramer labelling in vitro, SA-Fluor was used at the same final concentration served as a negative control. For Annexin V labelling *in vitro*, Annexin V was added at the same concentration but without Annexin V binding buffer.<br>Plasma EVs from mice i.v. injected with SA-647 were used as a control for specificity of the C1-Tetramer *in vivo*, those were measured at the same settings in a separate experiment. |
| 3.7 Serial dilutions. | State whether serial dilutions were performed on samples and note the dilution range and manner of testing. The fluorescence and/or scatter signal intensity would ideally be reported in standard units (see Section 4.3, 4.4) but arbitrary units can also be used. This data is best reported by plotting the recorded number events/concentration over a set period of time at different sample dilution. The median fluorescence intensity at each of the dilutions should also ideally be plotted on the same or a separate plot. | Serial dilutions were performed in each experiment for the fully stained samples, serially diluted samples were measured for the same amount of time and settings as the original samples. Three to four two-fold dilutions were done using filtered PBS and correlation analysis was performed showing a linear correlation for all samples ($R^2$>0.8), Particle/mL measured are plotted and shown in the supplementary figures. |
| 3.8. Detergent treated EV-samples | State whether samples were detergent treated to assess lability. If utilized, state what detergent was used, the end concentration of the detergent, and what the results were of the lysis. | Addition of NP-40 at a final concentration of 2.5% (v/v) was used for detergent controls which were acquired at the same settings as the corresponding samples. Exemplary graphs showing the reductions in $SSC_{low}$/CFSE+ or $SSC_{low}$/CTV+ events are shown for all experiments in the supplementary figures. |
| 4.1 Trigger Channel(s) and Threshold(s). | The trigger channel(s) and threshold(s) used for event detection. Preferably, the fluorescence calibration (Section 4.3) and/or scatter calibration (Section 4.4) should be used in order to report the trigger | Based on unstained, single-stained and/or fluorescence-minus-one controls, gates for event detection were set. All fluorescence detection was triggered at full power (488 nm; 200 mW. 642nm; 150 mW. 405 nm; 120 mW. 561; 200 mW). Scatter laser power (785 nm; 25 mW) was determined by the resolution of submicron sized particles using the MegaMix FSC and SSC mix (BioCytex), gating onto differently sized bead populations can be found in the main figures. |

| | | |
|---|---|---|
| | channel(s) and threshold(s) in standardized units. | |
| 4.2 Flow Rate / Volumetric quantification. | State if the flow rate was quantified/validated and if so, report the result and how they were obtained. | For all samples, ImageStream$^{TM}$ was set at 'low speed, high sensitivity', ~0.4 µLof sample was measured in the time span of 60 seconds. |
| 4.3 Fluorescence Calibration. | State whether fluorescence calibration was implemented, and if so, report the materials and methods used, catalogue numbers, lot numbers, and supplied reference units for the standards. Fluorescence parameters may be reported in standardized units of MESF, ERF, or ABC beads. The type of regression used, and the resulting scatter plot of arbitrary data vs standard data for the reference particles should be supplied. | Calibration to standard units was not performed. |
| 4.4 Light Scatter Calibration. | State whether and how light scatter calibration was implemented. Light scatter parameters may be reported in standardized units of nm2, along with information required to reproduce the model. | To test for submicron detection of fluorescent particles we used commercially available polystyrene beads (MegaMix FSC and SSC mix (BioCytex)). Using the side-scatter and the 488 nM laser we could discriminate beads in a range of 100-900 nM, highlighting the suitability of IFC to analyze heterogenous submicron-sized populations. We did not perform light-scatter calibration as we did not intend to use IFC for size approximation. |
| 5.1 EV diameter/surface area/volume approximation. | State whether and how EV diameter, surface area, and/or volume has been calculated using FC measurements. | For our study, sizing EVs by IFC was not relevant. We did examine average particle size for all our samples using Nanoparticle Tracking Analysis (NTA) which revealed an average EV size of ~131-152 nm. |

| | | |
|---|---|---|
| 5.2 EV refractive index approximation. | State whether the EV refractive index has been approximated and how this was done. | The EV refractive index has not been determined in this study. |
| 5.3 EV epitope number approximation. | State whether EV epitope number has been approximated, and if so, how it was approximated. | Epitope number has not been approximated using IFC. |
| 6.1 Completion of MIFlowCyt checklist. | Complete MIFlowCyt checklist criteria 1 to 4 using the MIFlowCyt guidelines. Template found at www.evflowcytometry.org. | See Suppl. Table 2. |
| 6.2 Calibrated channel detection range | If fluorescence or scatter calibration has been carried out, authors should state whether the upper and lower limits of a calibrated detection channel were calculated in standardized units. This can be done by converting the arbitrary unit scale to a calibrated scaled, as discussed in Section 4.3 and 4.4, and providing the highest unit on this scale and the lowest detectable unit above the unstained population. The lowest unit at which a population is deemed 'positive' can be determined a variety of ways, including reporting the 99th percentile measurement unit of the unstained population for fluorescence. The chosen method for determining at what unit an event was deemed | N/A |

| | | |
|---|---|---|
| | positive should be clearly outlined. | |
| 6.3 EV number/concentration. | State whether EV number/concentration has been reported. If calculated, it is preferable to report EV number/concentration in a standardized manner, stating the number/concentration between a set detection range. | Particle concentration of all samples was measured by Nanoparticle tracking analysis. Samples were then adjusted to $2\text{-}5^{10}$/mL prior to staining and IFC measurements. Detected concentrations of fluorescent EVs by IFC are shown in the serial dilution plots in the supplementary figures. |
| 6.4 EV brightness. | When applicable, state the method by which the brightness of EVs is reported in standardized units of scatter and/or fluorescence. | N/A |
| 7.1. Sharing of data to a public repository. | Provide a link to the experimental data in a public data repository. | IFC files can be obtained by contacting the corresponding author. |

**Supplementary Table S2** – Checklist representing the Minimal Information about a Flow Cytometry (FC) experiment for EV-FC-specific reporting (MIFlowCyt-EV checklist), as recommended by the Minimum Information for Studies of EVs (MISEV).

| **Requirement** | **Please Include Requested Information** |
|---|---|
| 1.1. Purpose | Determine the frequency of PS among blood EVs and analyze circulating and cell-bound PS+ EVs to investigate their clearance *in vivo* |
| 1.2. Keywords | Phosphatidylserine; Extracellular Vesicles; Imaging flow cytometry; superresolution microscopy; clearance |
| 1.3. Experiment variables | Experimental variables: a) murine plasma/serum from 6-8 weeks old female C57BL/6J mice were stained with CFDA-SE, CellTrace Violet, CD9, CD81, CD41, ApoE, C1-Tetramer (MFG-E8 derivative), Annexin V and/or MFG-E8-eGFP and measured by Imaging Flow Cytometry (and superresolution microscopy). b) Human plasma EVs from 5 healthy individuals were stained with CFDA-SE, CD9 and C1-Tetramer (MFG-E8 derivative) and measured by Imaging Flow Cytometry (and superresolution microscopy). |
| 1.4. Organization name and address | Biomedical Center Munich (LMU Munich)<br>Institute for Immunology<br>Grosshaderner Str. 9<br>82152 Planegg<br>Germany |
| 1.5. Primary contact name and email address | Dr. Jan Kranich |
| 1.6. Date or time period of experiment | August 2023-October 2024 |
| 1.7. Conclusions | PS-staining reagents have different sensitivities which is of particular importance when studying low PS frequency particles such as blood EVs in contrast to e.g. apoptotic cells. Moreover, PS represent a predominant marker of EVs in the blood and using the co-factor independent C1-Tetramer for PS-labelling, endogenous EVs can be stained *in vivo*. |
| 1.8. Quality control measures | The ImageStream™ calibration tool ASSIST® was used upon each startup to optimize performance and ensure consistency among experiments. Additionally, commercially available FITC-fluorescent polystyrene beads of distinct submicron sizes (Megamix-Plus FSC – 900, 500, 300 and 100 nm, and Megamix-Plus SSC – 500, 240, 200, 160 nm). |
| 2.1.1.1. (2.1.2.1., 2.1.3.1.) Sample description | All mice were housed under specific pathogen-free conditions at the Core Facility Animal Models of the Biomedical Center (Ludwig-Maximilians-University, Munich). All the procedures and animal housing conditions were approved by the government of Oberbayern. Blood from 6-8 weeks old female C57BL/6JNRj mice (Janvier) was collected by heart puncture. For serum, blood samples were allowed to coagulate for 30 min at RT whereas heparin was added to plasma samples. Subsequently, blood was centrifuged at 1,000 g for 10 min at 4°C and serum or plasma was diluted 1:1 with 1X cOmplete™ protease inhibitor cocktail (Roche). Alternatively, aliquots of plasma or serum were snap-frozen and stored at -20°C at this step. Samples were centrifuged sequentially at 2,000 g at 4°C and 10,000 g at RT for 10 min. Afterwards, 800-900 µL of sample were applied onto SEC columns (qEV1, Gen2 Izon) and |

| | |
|---|---|
| | EV-containing fractions were collected and concentrated to 300 µL. EV samples were not stored prior to the measurement but used freshly. Human plasma was collected from 5 healthy individuals (3 males, 2 females – age 21-60) into S-Monovette® Natrium Heparin tubes and processed as described above for murine samples. |
| 2.1.1.2. Biological sample source description | See above |
| 2.1.1.3. Biological sample source organism description | Mice - C57BL/6JNRj purchased from Janvier, females, 6-8 weeks old.<br>Humans – Healthy individuals, 3 males, 2 females, age 21-60 |
| 2.1.2.2. Environmental sample location | N/A |
| 2.3. Sample treatment description | Plasma or serum EVs were isolated as described above.<br>For IFC staining, all staining reagents were centrifuged at 18,000g for 30 min at 4°C. 1 µL of CFSE or CellTrace Violet were added to 40 µL of sample material to reach a final concentration of 10 µM. After addition of the membrane dyes, samples were incubated for 1 h at RT protected from light. Antibodies or staining reagents were added (or from predilutions) to the samples to reach the final concentrations as stated above in a total volume of 60 µL (to reach 60 µL among all samples spare volumes of filtered PBS were added if necessary). For Annexin V staining, 6 µL of 10X Annexin binding buffer were added to the samples while the total volume of 60 µL remained the same among all samples. Subsequently, staining was done for another hour at RT. Afterwards, samples were split in half and 2.5% (v/v) NP-40 was added to detergent controls whereas the same volume of PBS was added to the samples. Prior to IFC measurements, samples were diluted to 120 µL with filtered PBS or 120 µL filtered PBS with 1X Annexin V binding buffer. Aggregate controls contained the same final concentration of antibodies, staining reagents or membrane dyes diluted in filtered PBS. |
| 2.4. Fluorescence reagent(s) description | **Anti-mouse mAB:**<br>• Anti-mouse CD9 eFluor450 (clone: KMC8. isotype: rat IgG2a) (eBiosciences) (final concentration for staining in 60 µL: 120 ng/test)<br>• Anti-mouse CD9 APC/Fire$^{TM}$ 750 (clone: MZ3. isotype: rat IgG2a) (Biolegend) (final concentration for staining in 60 µL: 120 ng/test)<br>• Purified anti-mouse CD81 (clone: Eat-2. Isotype: Armenian Hamster IgG1k) fluorescently labelled with Mix-n-Stain CF568 Antibody labelling kit (Sigma-Aldrich) (final concentration for staining in 60 µL: 300 ng/test)<br>• Purified anti-mouse CD41 (clone: MWReg30. Isotype: rat IgG1k) (Biolegend) fluorescently labelled with Mix-n-Stain CF647 antibody labelling kit (Sigma-Aldrich) (final concentration for staining in 60 µL: 300 ng/test)<br>• Purified anti-mouse ApoE (clone: 3B3C32. Isotype: rat IgG2a,k) (Biolegend) labelled with Mix-n-Stain CF568 antibody labelling kit (Sigma-Aldrich) (final concentration for staining in 60 µL: 300 ng/test) |

| | |
|---|---|
| | **Anti-human mAB:**<br>• Anti-human CD9 BV421 (clone: M-L13. Isotype: mouse IgG1k) (BD biosciences) (final concentration for staining in 60 µL: 120 ng/test)<br>**Membrane dyes:**<br>• Carboxyfluorescein succinimidyl ester (CFSE, eBiosciences) (final concentration in 60 µL: 10 µM)<br>• CellTrace Violet (ThermoFisher Scienitific) (final concentration in 60 µL: 10 µM)<br>**PS-binding reagents:**<br>• C1-Tetramer (MFG-E8 biotinylated monomeric C1-domain teramerized with streptavidin) (unless stated otherwise final concentration was 5 nM in 60 µL)<br>• Annexin V FITC or AF647 (Biolegend) (final concentration varied from 0.005-830 nM in 60 µL)<br>• Recombinant full-length MFG-E8-eGFP (Kranich *et al.*, 2020)(final concentration varied from 0.005-830 nM in 60 µL)<br>**Anti-mouse mAB matched isotype controls:**<br>• For anti-mouse CD9 eFluor450 (eBiosciences) the following isotype control antibody was used: Rat IgG2a,l isotype control antibody BV421 (clone: RTK2759, Biolegend) (final concentration for staining in 60 µL: 120 ng/test)<br>• For anti-mouse CD9 APC/Fire$^{TM}$ 750 (clone: MZ3. isotype: rat IgG2a) the following isotype control antibody was used:<br>Rat IgG2a,l isotype control antibody APC-Cy7 (clone: RTK2758, Biolegend) (final concentration for staining in 60 µL: 120 ng/test)<br>• For purified anti-mouse CD81 (clone: Eat-2. Isotype: Armenian Hamster IgG1k) fluorescently labelled with Mix-n-Stain CF568 Antibody labelling kit (Sigma-Aldrich) the following isotype control antibody was used:<br>Purified armenian hamster IgG isotype ctrl antibody (clone: HTK888, Biolegend) fluorescently labelled with Mix-n-Stain CF568 Antibody labelling kit (Sigma-Aldrich) (final concentration for staining in 60 µL: 300 ng/test)<br>• For purified anti-mouse CD41 (clone: MWReg30. Isotype: rat IgG1k) (Biolegend) fluorescently labelled with Mix-n-Stain CF647 antibody labelling kit (Sigma-Aldrich) the following isotype control antibody was used: |

| | |
|---|---|
| | Purified rat IgG1k isotype ctrl antibody (clone: RTK2071, Biolegend) fluorescently labelled with Mix-n-Stain CF647 antibody labelling kit (Sigma-Aldrich) (final concentration for staining in 60 µL: 300 ng/test)<br>• For purified anti-mouse ApoE (clone: 3B3C32. Isotype: rat IgG2a,k) (Biolegend) labelled with Mix-n-Stain CF568 antibody labelling kit (Sigma-Aldrich) the following isotype control antibody was used:<br>Rat IgG2a,k isotype control antibody PE (clone:RTK2758) (final concentration for staining in 60 µL: 300 ng/test)<br>**Anti-human mAB matched isotype controls:**<br>• For Anti-human CD9 BV421 (clone: M-L13. Isotype: mouse IgG1k) (BD biosciences) the following isotype control antibody was used: mouse IgG1,k isotype control antibody BV421 (clone: MOPC-21, Biolegend) (final concentration for staining in 60 µL: 120 ng/test) |
| 3.1. Instrument manufacturer | Cytek |
| 3.2. Instrument model | ImageStream$^X$ MKII |
| 3.3. Instrument configuration and settings | The instrument is equipped with a 405nm, 488nm, 561nm, 592nm, 642nm, and 786nm lasers, three objectives (20x/40x/60x) and 2 CCD cameras. All data were acquired using the 60x objective with fluidics settings set to "low speed/high sensitivity". |
| 4.1. List-mode data files | IFC files can be obtained by contacting the corresponding author. |
| 4.2. Compensation description | For compensation, single stained controls were prepared for each fluorophore with BD CompBeads (Anti-Mouse Ig or Anti-Hamster) and BD CompBeads Negative controls (BD Biosciences). The compensation beads were acquired at the same laser settings and antibodies were diluted as stated above for the staining. Compensation matrices were calculated using the IDEAS software. |
| 4.3. Data transformation details | N/A |
| 4.4.1. Gate description | For gating, fluorescence-minus-one controls, unstained EVs and/or isotype controls were measured at the same settings and during each experiment. For C1-Tetramer labelling *in vitro*, SA-AF647 was used at the same final concentration served as a negative control. For Annexin V labelling *in vitro*, Annexin V was added at the same concentration but without Annexin V binding buffer. |
| 4.4.2. Gate statistics | Particles/mL, Particles/gate, frequency of parent (%) |
| 4.4.3. Gate boundaries | See above |

**Supplementary Table S3** – Reagents

| **Vendor** | **Cat#** | **Reagent** | **Clone/reference** |
|---|---|---|---|
| ThermoFisher | C34557 | CellTrace™ Violet | |
| eBioscience | 65-0850-84 | Carboxyfluorescein succiminidyl ester (CFSE) | |
| Biolegend | 405237 | Streptavidin AF647 | |
| Biolegend | 405235 | Streptavidin AF488 | |
| Biolegend | 405225 | Streptavidin BV421 | |
| Biotium | 29035 | Streptavidin CF568 | |
| Aurion | 806.099 | Streptavidin-gold (6 nm) | |
| Biolegend | 64090 | Annexin V FITC | |
| Biolegend | 64091 | Annexin V AF647 | |
| Biolegend | 674802 | ApoE (unlabelled) | 3B3C32 |
| Biolegend | 13390 | CD41(unlabelled) | MWReg30 |
| Biolegend | 10130 | Fc-block (mouse) | 93 |
| Miltenyi | 130-059-901 | Fc-block (human) | |
| Biolegend | 40040 | Rat IgG1, k isotype ctrl (unlabeled) | RTK2071 |
| Biolegend | 40050 | Rat IgG2a, k isotype ctrl PE | RTK2758 |
| Biolegend | 40052 | Rat IgG2a, k isotype ctrl APCCy7 | RTK2758 |
| Biolegend | 40053 | Rat IgG2a, k isotype ctrl BV421 | RTK2758 |
| Biolegend | 40090 | Armenian Hamster IgG isotype ctrl (unlabeled) | HTK888 |
| Biolegend | 124802 | CD9 (unlabeled) | MZ3 |
| Biolegend | 104901 | CD81 (unlabeled) | Eat-2 |
| Biolegend | 104903 | CD81 biotinylated | Eat-2 |
| eBiosciences | 14-0091-82 | CD9 (unlabeled) | eBioKMC8 (KMC8) |
| eBiosciences | 48-0091-82 | CD9 eFluor450 | eBioKMC8 (KMC8) |
| Biolegend | 12481 | CD9 APC/FireTM 750 | MZ3 |
| CellSignaling | 49295 | Albumin (unlabeled) | polyclonal |
| Biolegend | 312102 | CD9 (unlabeled) | HI9a |
| eBiosciences | 13-0098-82 | CD9 biotinylated | eBioSN4 |
| Bethyl-Fortis | A80-192P | Albumin-HRP | polyclonal |
| Jackson ImmunoResearch | 711-035-152 | Anti-rabbit IgG-HRP | |
| Jackson ImmunoResearch | 715-035-150 | Anti-mouse IgG-HRP | |
| CellSignaling | 7707 | Anti-rat IgG-HRP | |
| Biolegend | 10122 | CD11b-APCCy7 | M1/70 |

| | | | |
|---|---|---|---|
| Biolegend | 11550 | CD19-FITC | 6D5 |
| ThermoFisher | L34955 | LIVE/DEAD$^{TM}$ fixable violet dead cell stain kit | |
| Biolegend | 10022 | CD3 BV421 | 17A2 |
| ThermoFisher | | CD19 eFluor450 | |
| Biolegend | 11732 | CD11c BV421 | N418 |
| self-made | | MFG-E8-eGFP | (Kranich et al., 2020) |
| self-made | | MFG-E8 biotinylated C1 monomer | Patent publication No. US 2024/0125806 A1 |
| Biolegend | 480157 | MFG-E8 biotinylated C1 monomer (ApoMonomer) | |
| Sigma-Aldrich | MX488AS100 | Mix-n-Stain$^{TM}$ CFTM488 labeling kit | |
| Sigma-Aldrich | MX568S100 | Mix-n-Stain$^{TM}$ CFTM568 labeling kit | |
